# A method for identifying environmental stimuli and genes responsible for genotype-by-environment interactions from a large-scale multi-environment data set

**DOI:** 10.1101/2021.10.25.465681

**Authors:** Akio Onogi, Daisuke Sekine, Akito Kaga, Satoshi Nakano, Tetsuya Yamada, Jianming Yu, Seishi Ninomiya

## Abstract

It has not been fully understood in real fields what environment stimuli cause the genotype-by-environment (G × E) interactions, when they occur, and what genes react to them. Large-scale multi-environment data sets are attractive data sources for these purposes because they potentially experienced various environmental conditions. Here we developed a data-driven approach termed <u>E</u>nvironmental <u>C</u>ovariate Search Affecting <u>G</u>enetic <u>C</u>orrelations (ECGC) to identify environmental stimuli and genes responsible for the G × E interactions from large-scale multi-environment data sets. ECGC was applied to a soybean (*Glycine max*) data set that consisted of 25,158 records collected at 52 environments. ECGC illustrated what meteorological factors shaped the G × E interactions in six traits including yield, flowering time, and protein content and when they were involved. For example, it illustrated the relevance of precipitation around sowing dates and hours of sunshine just before maturity to the interactions observed for yield. Moreover, genome-wide association mapping on the sensitivities to the identified stimuli discovered candidate and known genes responsible for the G × E interactions. Our results demonstrate the capability of data-driven approaches to bring novel insights on the G × E interactions observed in fields.

**Key message:** The proposed method is able to identify environmental stimuli and genes responsible for the G × E interactions observed in multi-environmental trials. The method is based on similarity search between genetic correlation and environmental stimuli among environments.

## Introduction

Genotype-by-environment (G × E) interactions have been one of main interests in plant research for decades (Mather & Jones, 1958; van Eeuwijk *et al*., 2005; Des Marais *et al*., 2013). However, understanding the interactions in fields is not an easy task because a number of players including environmental stimuli and genes can be involved in at various growth stages. Thus, it has been a great challenge to depict comprehensive landscapes on how genes and environment stimuli cause the G × E interactions together in fields. So far, studies have successfully used environmental stimuli in statistical models to map quantitative trait loci (QTLs) and/or predict crop phenotypes (Malosetti *et al*., 2013; Jarquin *et al*., 2014; Li *et al*., 2018; Millet *et al*., 2019; Guo *et al*., 2020). These studies show the usefulness of a reaction norm approach where phenotypes/genotypic values are regressed on quantitative indices of environment stimuli to model the sensitivity of genotypes. An important point of this approach is, however, that the sensitivity of genotypes is not necessary associated with the observed G × E interactions. Methods to identify environmental stimuli and genes directly related to the G × E interactions have been lacked.

Here, we propose a novel method termed <u>E</u>nvironmental <u>C</u>ovariate Search Affecting <u>G</u>enetic <u>C</u>orrelations (ECGC) to reveal quantitative environmental stimuli (referred to as environmental covariates) and genetic architecture underpinning the G × E interactions using large-scale multi-environment data sets. ECGC searches environmental covariates whose similarity matrices between environments are significantly correlated with genetic correlation matrices between environments which can be regarded as indicators of the G × E interactions (Hayes *et al*., 2016). This proposed method is able to identify the environmental stimuli and genes directly related to the observed G × E interactions. Moreover, because genetic correlations between environments can be estimated using mixed models, ECGC is robust to missing records and unbalanced data structure that often characterize multi-environment data. Here we applied the proposed method to a large-scale multi-environment data of soybean (*Glycine max*). The traits were days to flowering (DTF), days to maturity (DTM), stem length (SL, cm), protein content of seeds (PR, %), yield (YI, kg/a) and seed weight (SW, g/100 seeds).

## Materials and Methods

### Model description

As described above, ECGC searches environmental covariates whose similarity matrices between environments are correlated with genetic correlation matrices between environments. Here it will be illustrated how the similarity of environmental covariates is associated with genetic correlation between environments.

For the *j*^th^ environment, the phenotype adjusted for variations due to years and management conditions, 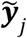, is decomposed as

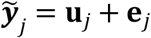

where **u**_*j*_ and **e**_*j*_ are the additive genetic effect and residual, respectively. Note that 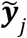 and **u**_*j*_ include all adjusted phenotypes and additive genetic effects of genotypes evaluated at the environment. **u**_*j*_ and **e**_*j*_ are assumed to follow multivariate normal distributions (MVN) as 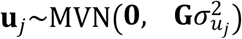 and 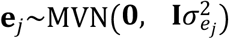, respectively, where **G** is the genomic relationship matrix and **I** is the identity matrix. After scaling with the genetic standard deviation 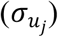, the additive genetic effect is assumed to be further decomposed into two components, one affected by an environmental covariate, *x*_*j*_, and the other is free of it, as

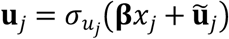

where **β** is the slopes of varieties that represent the sensitivity to the environmental covariate, and 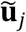 is the residual genetic effects. Here **β** is assumed to be 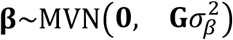 which means 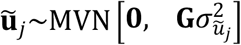 where 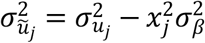. Then the genetic covariance of the additive genetic effects between environments *j* and *k* can be represented as

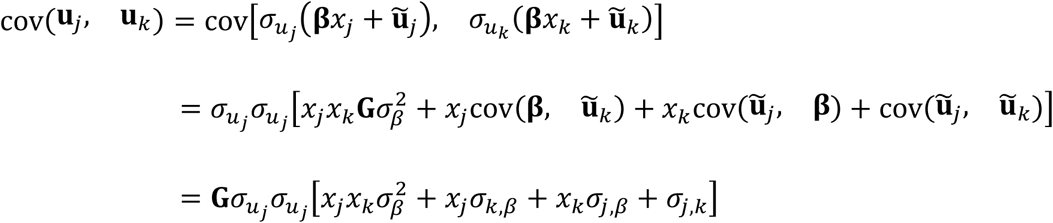

by letting 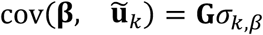 and 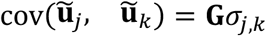. Thus, the genetic covariance between environments *j* and *k*, 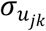, is

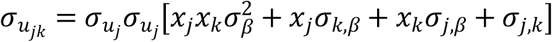

and the genetic correlation between these environments, *P*_*j*,*k*_, is

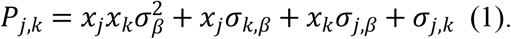

Eq. 1 illustrates how the similarity of environmental covariates (*x*_*j*_*x*_*k*_) is related to genetic correlation between environments (*P*_*j*,*k*_). Genetic correlations between environments poses information of relative magnitudes of genotypic values among varieties that change across environments, i.e., G × E interactions. Thus, ECGC searches the environment covariates that are associated with the G × E interactions by scanning cor_*j*,*k*_(*P*_*j*,*k*_, *x*_*j*_*x*_*k*_) where cor_*j*,*k*_ means taking correlations across all combinations of environments. The squared cor_*j*,*k*_(*P*_*j*,*k*_, *x*_*j*_*x*_*k*_) (i.e., *r^2^*) represents the proportion of the variance of 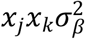 in the variance of *P*_*j*,*k*_. The significantly detected environmental covariates will be able to be used for various purposes. Here we focus on revealing genes underlying the G × E interactions. To this end, GWA mapping was conducted on the slopes (**β**) associated with the detected environmental covariates as the final step of ECGC.

Eq. 1 shows that environmental covariates (*x*_*j*_*x*_*k*_) are associated with G × E interactions (*P*_*j*,*k*_) when 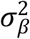 is not zero and the term *x*_*j*_*σ*_*k*,*β*_ + *x*_*k*_*σ*_*j*,*β*_ + *σ*_*j*,*k*_ does not conceal 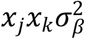. This fact implies that 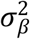 itself is not an evidence that the environmental covariate is associated with G × E interactions; even when 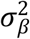 significantly deviates from zero, the effect of 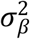 on the genetic correlation can be concealed by the term *x*_*j*_*σ*_*k*,*β*_ + *x*_*k*_*σ*_*j*,*β*_ + *σ*_*j*,*k*_. That is, non-zero 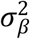 is a necessity condition for the association rather than a sufficient condition. On the other hand, significant correlations between *P*_*j*,*k*_ and *x*_*j*_*x*_*k*_ can be the direct evidence of association between the environmental covariate and G × E interactions because, when the term *x*_*j*_*σ*_*k*,*β*_ + *x*_*k*_*σ*_*j*,*β*_ + *σ*_*j*,*k*_ conceals 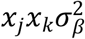, *x*_*j*_*x*_*k*_ is no longer correlated with *P*_*j*,*k*_. This fact also implies that statistical testing on 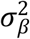 is not an alternative of ECGC. Both approaches (ECGC and testing on 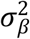) have different roles (detecting environment covariates associated with G × E interactions and genotype sensitivity, respectively).

### Data analysis overview

To apply ECGC to real data, five steps are required.

1. Calculation of environmental covariates (*x*_*j*_).
2. Calculation of similarities of environmental covariates between environments (*x*_*j*_*x*_*k*_).
3. Estimation of genetic correlation between environments (*P*_*j*,*k*_).
4. Scanning environmental covariates associated with genetic correlation based on cor_*j*,*k*_(*P*_*j*,*k*_, *x_j_x_k_*).
5. Estimation of slopes (**β**) which representing the sensitivity of varieties to the environmental covariates detected in Step 4 and conduct GWA mapping on the slopes.

These steps correspond with Fig. 1b–f. In the following sections whose titles start as “ECGC step”, we explain how these steps were conducted in our analyses using a large-scale multi-environment data of soybean. Preceding to these steps, we also conducted model development and fitting for each environment-trait combination to calculate the phenotypes adjusted with fixed effects 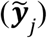. This step may not be always necessary for ECGC, but required for our data. This step is referred to as Step 0.

**Fig. 1.**
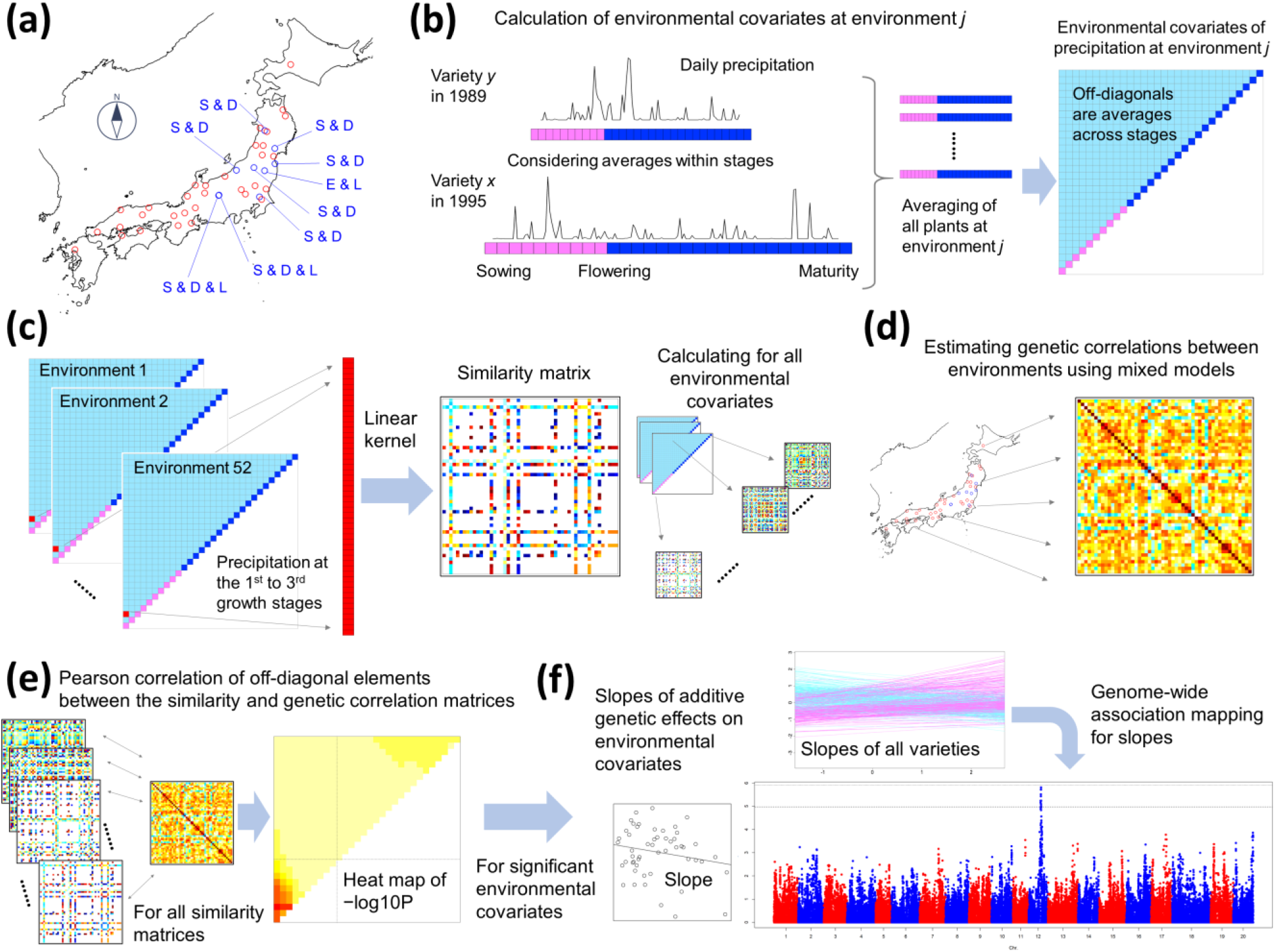
Illustration of the framework of ECGC. **(a)** Forty-one fields and 52 environments were analysed in this study. The blue points indicate fields where multiple management conditions were regarded as different environments. E and L denote early and late sowing, and D and S denote dense and sparse plant densities, respectively. **(b)** Calculation of environmental covariates representing each environment. Precipitation is illustrated as a meteorological factor. The whole growth periods of plants were divided into 30 stages (10 from sowing to flowering and 20 from flowering to maturity), and the meteorological values within each stage were averaged. These averaged values were again averaged across all plants at the environment, thus creating 30 environmental covariates (red and blue boxes in the upper-right triangle). The environmental covariates were then averaged across all spans within the 30 stages (1^st^ to 2^nd^ stages, 1^st^ to 3^rd^ stages, etc.), thus generating 435 additional covariates (pale-blue boxes in the triangle). In total, 465 (30 + 435) environmental covariates for each trait-meteorological factor combination were generated. Because 14 meteorological factors were considered, 465 × 14 = 6510 environmental covariates were generated for each trait. **(c)** Calculation of the similarity matrix of environmental covariates between environments. The figure illustrates the similarity matrix of precipitation at the 1^st^ to 3^rd^ growth stage, as an example. For each environmental covariate, values were extracted from the 52 environments and a 52 × 52 similarity matrix was calculated using the linear kernel. **(d)** Estimation of genetic correlations between environments. Using mixed models and genome-wide SNPs, genetic correlations were estimated for the 52 environments in a pairwise manner. **(e)** Calculation of the Pearson correlation coefficient (*r*) of off-diagonal elements between the similarity and genetic correlation matrices. *r* was calculated for all similarity matrices, and *−*log10 *P* values are presented as heat maps. **(f)** Genome-wide association mapping for uncovering genetic architecture. For the environmental covariates significantly detected in (E), slopes were estimated for each variety by regressing the additive genetic effects estimated using mixed models on the environmental covariates. Genome-wide association mapping was conducted on the slopes.

### Multi-environment data of soybean

The data set used in this study consisted of 25,158 records of 624 varieties evaluated at 41 fields (Supplementary Data 1). This data set was extracted as described below from a historical data of soybean including 72,829 records of 6,106 varieties evaluated at 440 fields over 55 years in Japan (from 1961 to 2015). The overview of this historical data is provided in Supplementary Methods 1. In nine fields, multiple management conditions (e.g. early or late sowing dates) were consistently conducted in multiple years. For these fields, different management conditions were defined as different environments as described later. As a result, a total of 52 environments were defined from 41 fields (Table S2, Fig. 1a). Thus, in this study, “environments” denote the combinations of fields (locations) and management conditions.

### DNA extraction and SNP genotyping

Seeds of varieties included in the historical data were collected from the breeding centres and NARO genebank (as many seeds as possible). The seeds were sown in pots and grown until the first trifoliate leaves emerged in greenhouses. Total genomic DNA was extracted from the first trifoliate leaves of one plant using a method based on guanidine hydrochloride and proteinase K (Khosla *et al*., 1999), with modifications. Among the ~2,000 varieties with extracted DNA, 573 varieties were genotyped using the Axiom SNP custom array (ThermoFisher, MA, USA), which was designed for Japanese soybean varieties. These 573 varieties consisted mainly of breeding lines that had been evaluated until later generations (typically F8 or later) as promising lines. In addition, from the ~2,000 varieties, 149 varieties that were registered as major varieties by the Ministry of Agriculture, Forestry and Fisheries and 187 varieties included in the soybean core collection (Kaga *et al*., 2012) were also genotyped using the array. The 573 varieties were selected to avoid overlap with the 336 (149 + 187) varieties. Among the SNPs genotyped, SNPs that showed high genotyping quality (i.e. SNPs termed ‘PolyHighResolution’ in the Axiom genotyping system) and could be mapped to the Williams 82 genome assembly version 2.0 (Wm82.a2.v1/Glyma 2.0) were extracted (138,555 SNPs) (Table S1). The average call rate of SNPs was 0.998 (SD ± 0.003). Missing genotypes were imputed using Beagle 4.1 (08Jun17.d8b) (Browning & Browning, 2016).

### Extraction of phenotypic records

The varieties that had SNP genotypes from the Axiom custom array were subjected to subsequent analyses. Because the names of varieties usually change with the advancement of generations, the variety names were integrated using the names at the last generations. Among the 440 fields included in the historical data, 41 fields were selected for analyses because of the number of records and balanced geological locations across Japan (Fig. 1a). For each field, phenotypic values that were out of the mean ± 3SD (standard deviation) range were treated as missing values.

The following six traits were selected for analyses: DTF (days), DTM (days), SL (cm), PR (%), YI (kg/a) and SW (g/100 seeds). These traits were selected because of their importance for breeding and the number of phenotyped records. In Japanese soybean breeding, to unify the measuring methods of traits, regulations on trait measurement were established in 1954 by a committee where the Ministry of Agriculture and Forestry at the time took the leading role. Although these regulations seem to have undergone several minor updates to date, trait definitions were largely common across the study period. Briefly, DTF was defined as the date when 40%–50% of the buds of the strains reached flowering. DTM was the date when 80%–90% of the pods of the strains showed variety-unique colours at maturity. SL was the length of the main culm between the ground surface and the growth point. PR was mainly measured using near-infrared spectroscopy, but there seemed to be some variations in the measuring methods and used machines. YI and SW were defined as the weight with 15% moisture, although there also seemed to be some variations in the moisture percentage among stations and years.

Records that had at least one observation for these six traits were extracted for subsequent analyses. As a result, the extracted records consisted of 25,158 records of 624 varieties evaluated at 41 fields (Supplementary Data 1). The summaries of the extracted phenotypic data and used varieties are presented in Tables S2 and S3, respectively.

### Meteorological factors

Meteorological information was taken from MeteoCropDB ver. 1.0 (Kuwagata *et al*., 2011). The database was designed to help the utilisation of crop growth models and provides daily values at each weather station of the Japan Meteorological Agency (JMA). Meteorological information for each environment was taken from the nearest JMA station. In this study, 13 meteorological factors, i.e. mean temperature (°C), maximum temperature (°C), minimum temperature (°C), precipitation (mm), vapour pressure (hPa), vapour pressure deficit (hPa), relative humidity (%), minimum relative humidity (%), wind speed (m/s), maximum wind speed (m/s), hours of sunshine (h), solar radiation (W/m^2^) and potential evapotranspiration (mm), were used. Among these factors, wind speed and maximum wind speed were adjusted as observed at 2.5 m of altitude by the developer of the database from the original values of the JMA. Solar radiation was estimated by the database developer from the observations of the JMA regarding hours of sunshine. Potential evapotranspiration was also estimated by the developer based on mean temperature, vapour pressure, wind speed, solar radiation and air pressure. The values of the other factors were according to the observations of the JMA. In addition to these factors, photoperiod (h) was calculated for each environment. For simplicity, daily length was also considered as a meteorological factor. The values of these meteorological factors are included in Supplementary Data 1, together with the phenotype records.

### ECGC Step 0: Developing mixed models at each environment

Single-trait mixed models were fitted to the records at each environment. This process had two purposes; the first was to eliminate variations attributable to years and management conditions from phenotypic values, that is, to create 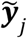. The adjusted phenotypes 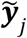 were then used for estimating genetic correlations between environments (ECGC Step 3). The second was to estimate the additive genetic effects of varieties at each environment. The additive genetic effects were then used to estimate the slopes of genetic effects on environmental covariates (ECGC Step 5).

The year effects were added as fixed effects in the mixed models. The conditions of three management methods (plant density, fertiliser and sowing date) were modelled in various ways. In nine fields (F04, F07, F08, F09, F22, F24, F25, F27 and F34), multiple management conditions were consistently conducted in multiple years. For these fields, different conditions were defined as different environments resulting in a total of 52 environments (Table S2, Fig. 1a).

In most fields, however, the management conditions often varied across years. When the effects of management conditions were indistinguishable from those of year effects, the effects were absorbed to the year effects and not modelled. Otherwise, the effects were added in the mixed models. The modelling schemes varied according to the environments and management methods. Although the conditions of plant density, fertilizer, and sowing date are intrinsically continuous variables, when the number of conditions at an environment was few, these conditions were regarded as categorical variables and the effects were modelled as fixed effects. Otherwise, the conditions were regarded as continuous and the effects were model using basis functions (e.g. B-splines, Hastie *et al*., 2009). In each case, the interactions between genotypes and management were also considered. Variance components were estimated using REML implemented in airemlf90 (Misztal *et al*., 2002), and model selection was conducted using the AICs provided by the programme. The covariance structure of additive genetic effects among varieties (i.e. genomic relationship matrix) was defined using the genome-wide SNPs (VanRaden, 2008). The A.mat function provided by the R package, rrBLUP (ver. 4.6.1) (Endelman, 2011; R Core Team, 2020), was used for calculation (Supplementary Data 2). Thus, the mixed models applied here were the so-called genomic or genome-enabled BLUP (GBLUP) (de los Campos *et al*., 2013). The details of this procedure are described for each environment in the Supplementary Methods 2. Narrow-sense SNP heritability estimated by the selected models are presented in Table S4.

### ECGC Step 1: Calculation of environmental covariates

Environmental covariates were calculated for each environment, i.e., combination of fields and management conditions. Because trials had been conducted for multiple years at each field (Table S1), in principle, environmental covariates were calculated by averaging meteorological values across years. The details are explained below.

Even on the same calendar day, the growth stages of plants can vary depending on the sowing date and growth speed, which depend on the environmental conditions (e.g. temperature) and genotypes. Thus, a meteorological event (e.g. high or low temperature) on a calendar day can have different effects on plants with different growth stages. To consider the difference in growth stage, the growth period between the sowing dates and days of flowering was divided into 10 equal-sized stages (on average 5.5 ± 1.2 days per stage), and the period between the days of flowering and days of maturity was divided into 20 equal-sized stages (3.8 ± 0.5 days per stage). For each variety at each environment and year, the daily meteorological values within stages were averaged (Fig. 1b).

Meteorological values representing each environment were then obtained by averaging the above-mentioned meteorological values of all plants included in the environment, yielding 30 values for the 30 growth stages for each environment (Fig. 1b, Tables S5–10). In addition, meteorological values were averaged across stages, e.g. averages across the 1^st^ to 2^nd^ stages, across the 1^st^ to 3^rd^ stages, etc. This procedure was inspired by the joint genomic regression analysis (Li *et al*., 2018). As a result, 435 (30 × 29 / 2) additional representative values were generated, resulting in 465 values for each environment (Fig. 1b). That is, 465 × 14 (meteorological factors) = 6510 environmental covariates were generated.

Two issues were notable in this procedure. First, the number of stages will depend on the prior knowledge or assumptions on growth stages. For soybean, typically five and eight growth stages are assumed for the vegetative and reproductive phases (Fehr & Caviness, 1977). The number of stages used here (30) were determined to be able to discriminate these known growth stages. If a crop experiences more growth stages, setting a larger number will facilitate interpretation. Second, although the growth stages of plants can be represented using cumulative temperature and/or photoperiod, we avoided using these indices, to simplify the procedures as much as possible.

### ECGC Step 2: Calculation of similarities of environmental covariates between environments

For each environmental covariate (i.e. combination of growth stage and meteorological factor), 52 representative values corresponding to 52 environments were used to calculate the similarity between environments (Fig. 1c). The similarity was defined using a linear kernel. That is, the similarity between the *j*^th^ and *k*^th^ environments were calculated as *x_j_x_k_* where *x_j_* and *x_k_* indicate the deviations of the environmental covariate values at environments *j* and *k*, respectively. As a result, a 52 × 52 similarity matrix was obtained for each environmental covariate. Total 465 growth stages × 14 meteorological factors = 6510 similarity matrices were obtained for each trait.

### ECGC Step 3: Estimation of genetic correlation between environments

Genetic correlations between environments were estimated bivariate mixed models in a pairwise manner. The model can be written as

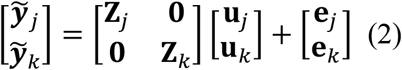

where **Z**_*j*_ and **Z**_*k*_ are the design matrices. Here it is assumed that 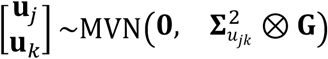 and 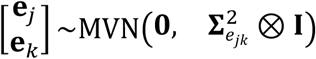 where 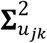 and 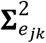 are the genetic and residual covariance matrices, respectively and ⊗ denotes the Kronecker product. To fit this model, first, the records at each environment (i.e., records in 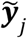 and 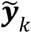) were matched to each other. Records were matched according to the varieties and evaluation years. That is, same varieties evaluated at the same year were matched to each other. When multiple records could be matched (i.e., when a variety was evaluated at an environment multiple times in a single year), matching was determined randomly. As a result of matching, the phenotypic data consisting of 25,158 records was arranged to a matrix of 7,887 by 52 (Supplementary Data 3 and 4). Note that most records did not match to any other records. Thus, the resulting 7,887 by 52 matrix was sparse: the non-missing proportion was at most 25,158 / (7,887 × 52) = 0.061. Because PR was absent in 22 environments, the matrix for PR becomes 7,887 by 30 (Supplementary Data 3 and 4).

Subsequently, for each trait, genetic correlations between environments were estimated in a pairwise manner. First, the bivariate mixed model (Eq. 2) was fitted to all pairs of environments (52 × 51 / 2 = 1326 pairs), to estimate the genetic covariances between environments 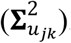. The diagonal elements of the covariance matrix (i.e. genetic variances) were estimated by averaging the estimates of these bivariate model analyses, which were duplicated 51 times. This pairwise manner of estimation of covariance matrices can be interpreted as a pseudo-likelihood-based approach (Fieuws & Verbeke, 2006). The resulting genetic covariance matrix was then set as a positive definite using the nearPD function of the Matrix package (ver. 1.2-18) of R, and a genetic correlation matrix was calculated by standardising the covariance matrix (Fig. 1d, Tables S11–16).

It is notable that genetic correlations between environment can be estimated using methods other than the pairwise estimation described here (e.g., standard multivariate models or factor analytic models). But the pairwise estimation will be one of the easiest methods to conduct in particular when the number of environments is great.

### ECGC Step 4: Scanning environmental covariates associated with the genetic correlation

Now we have the 6510 (465 growth stages × 14 meteorological factors) similarity matrices of environmental covariates and six (number of traits) genetic correlation matrices. The sizes of both kind matrices were 52 (number of environments) × 52 for the traits except for PR where the sizes were 30 × 30. The Pearson’s product moment correlation coefficients between the upper (or lower) triangle off-diagonals of the similarity matrices and genetic correlation matrices were then calculated (Fig. 1e) using the cor.test function of the R package stats (ver. 4.0.3). *P* values of the coefficients were also calculated using the function. The significance of the coefficients was judged after Bonferroni correction of the *P* values. Considering the number of environmental covariates (6510) and traits (6), the threshold of significance was set at 0.05 / (6510 × 6) = 1.28e–6.

Even after Bonferroni correction, 10397 environmental covariate/trait combinations remained significant. This was attributed to redundancy in the environmental covariates. For example, a value at the 10^th^ to 12^th^ growth stages was often highly correlated with a value at the 9^th^ to 13^th^ growth stages. To eliminate this redundancy, environmental covariates were clustered using the hierarchical clustering implemented in the hclust function of R. Similarities were defined based on Euclidean distance, and complete-linkage clustering was used. The number of clusters was determined using the gap statistic (Tibshirani *et al*., 2001). The clustering results are presented in Fig. S1. When multiple environmental covariates were significant at a cluster, the combination with the highest *r^2^* was selected, resulting in 555 significant environmental covariate/trait combinations (Table S17).

### ECGC Step 5: Estimation of slopes and GWA mapping on the slopes

For each significant environmental covariate/trait combination, the additive genetic effects of the 624 varieties at each environment were regressed on the environmental covariates, and the slopes (**β**) were calculated for each variety (Fig. 1f, Supplementary Methods 3, Table S18). The additive genetic effects were estimated using single-trait mixed models, as described in ‘ECGC Step 0: Developing mixed models at each environment’. The additive genetic effects were scaled with the additive genetic variance 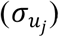, and the meteorological values were standardised before this regression analysis. Considering these slopes as phenotypic values, GWA mapping was conducted (Fig. 1f) using the GWAS function of the rrBLUP package (ver. 4.6.1). The genomic relationship matrix was calculated using the A.mat function of the package. No principal components were included in the models because *P* values did not inflate (Fig. S2). SNPs with minor allele frequencies > 0.05 were subjected to the statistical tests. This conservative threshold (0.05) was adopted to prevent false positives. This GWA mapping involved more than 6.40e+7 statistical tests (on average, 115,312 SNPs × 555 environmental covariate/trait combinations); thus, the application of strict multiple testing correction, such as Bonferroni correction, is unrealistic. Instead, we applied a method that controlled the false discovery rate (FDR) (Storey & Tibshirani, 2003). The threshold of significance was calculated using an R script implemented in the GWAS function of rrBLUP, with modifications. The threshold for FDR < 0.05 was 9.086278e−06 (5.041614 in the −log10p expression). The distribution of the *P* values is shown in Fig. S2. By grouping significant SNPs that were less than 100k bp apart from each other into the same regions, 1486 regions were detected for 270 environmental covariate/trait combinations with overlaps (Table S19).

To verify the validity of these associations, subsampling analyses were conducted for these 270 combinations. Specifically, 500 (80 %) varieties were randomly selected from the 624 varieties and subjected to GWA mapping. Random sampling was adopted because the aim of this subsampling is to perturbate the genetic structure which might produce false positives. This procedure was repeated 10 times. Then replications where the regions detected using the full data also showed significant associations were counted (threshold −log10p = 5.041614; Table S19). Regions with counts > 4 were regarded as reliable results. As a result, 948 regions were remained for 179 environmental covariate/trait combinations. These regions were found to be constituted with 39 regions by removing overlaps between combinations (Table S19).

Orthologs of *Arabidopsis thaliana* mapped in the detected regions were extracted from JBrowse provided by Phytozome ver.12.1 (https://phytozome.jgi.doe.gov/jbrowse/index.html?data=genomes/Gmax). Gene ontology of these orthologs were surveyed with overrepresentation analyses provided by PANTHER ver.16.0 (http://pantherdb.org/) using *Arabidopsis* genes as a reference.

### Simulation analyses

Simulation analyses were conducted to assess the performance to ECGC to detect environment covariates. As illustrated in Eq. 1, the detection power of ECGC depends on the proportion of the variance of 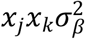 to that of *P*_*j*,*k*_ (i.e., *r^2^*) and the number of environments denoted as *M*. The detection power was expected to increase as *r^2^* and *M* increase. For the sake of simplification, *σ*_*j*,*β*_ = 0 and 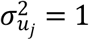 for any environment *j*, and 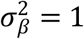 throughout the simulations. Consequently, *P*_*j*,*k*_ = *x*_*j*_*x*_*k*_ + *σ*_*j*,*k*_. The genetic correlation matrices between environments were simulated as follows.

1. Assign values for *r^2^* and *M* from grids 0.01, 0.018, 0.025, 0.035, 0.053, 0.075, 0.1, and 0.15, and 5, 10, 15, 20, 30, 40, 50, and 60, respectively.
2. Generate *x*_*j*_ from the standard normal distribution for all *j* (1 ≤ *j* ≤ *M*), and calculate the variance of *x*_*j*_*x*_*k*_ (1 ≤ *j* < *k* ≤ *M*).
3. Determine the variance of *σ*_*j*,*k*_ according to *r^2^* and the variance of *x*_*j*_*x*_*k*_.
4. Generate *σ*_*j*,*k*_ from the LKJ distribution using the R package rethinking (ver. 2.13) (McElreath, 2020). Parameter *η* of the distribution was arbitrary set to four. Scale *σ*_*j*,*k*_ according to the variance of *σ*_*j*,*k*_.
5. Generate a symmetric matrix by adding *x*_*j*_*x*_*k*_ to *σ*_*j*,*k*_. Here *x*_*j*_*x*_*j*_ (i.e., diagonal elements) is set to the average of *x*_*j*_*x*_*j*_ (1 ≤ *j* ≤ *M*).
6. Convert the symmetric matrix of (5) to the correlation matrix.

For each combination of *r^2^* and *M*, the genetic correlation matrix was simulated 2000 times. For each simulated matrix, 99 additional environment covariates that were not associated with the correlation matrix were also simulated as true negatives. Then the Pearson correlation between *P*_*j*,*k*_ and *x*_*j*_*x*_*k*_ was tested for these simulated environmental covariates. The performance of ECGC was assessed using the ROC curves drawn by the R package ROCR (ver. 1.0-11) (Sing *et al*., 2005).

## Results

The multi-environment data set used here consisted of 25,158 records of 624 varieties that were evaluated from 1961 to 2015 at 41 fields. The included varieties consisted of cultivars and breeding lines developed in Japan for the sake of food production, such as tofu (Table S3). Among the 41 fields, in nine fields, multiple management conditions (early/late sowing dates and sparse/dense plant densities) were consistently conducted in multiple years. For these fields, different management conditions were defined as different environments as described in Materials and Methods. As a result, a total of 52 environments were defined from 41 fields (Fig. 1a and Table S2).

The steps of ECGC are shown in Fig. 1b–f. In the first step, environmental covariates representing each environment were calculated (Fig. 1b). Here, we considered 14 meteorological factors as the environmental stimuli (Materials and Methods). We divided the whole growth period of a plant, from sowing to maturity, into 30 stages; this was achieved by dividing the growth period (from sowing to flowering) into 10 stages, and the period from flowering to maturity into 20 stages (Fig. 1b). For each meteorological factor (e.g. daily mean temperature), the meteorological values within each stage were averaged, yielding 30 values for each plant. These values were then averaged across plants evaluated at the environment for all the years, thus yielding 30 values for each environment (Tables S5–S10). Subsequently, the values were further averaged across the 30 stages, i.e. across the 1^st^ to 2^nd^ stages, the 1^st^ to 3^rd^ stages, etc. This procedure yielded 465 (30 + 30 × 29 / 2) environmental covariates for each trait–meteorological factor combination. Thus, total 465 × 14 (meteorological factors) = 6510 environmental covariates were generated.

In the second step, for each environmental covariate (e.g. precipitation at the 1^st^ to 3^rd^ stages), values were extracted from the 52 environments, and the linear kernel (i.e. the similarity matrix) between environments was calculated (Fig. 1c). In the third step, genetic correlations of additive genetic effects between environments were estimated using mixed models for each trait (Fig. 1d, Tables S11–S16, Fig. S3). Estimated genetic correlations suggest strong G × E interactions particularly for traits other than SW (Fig. S3). In the fourth step, the genetic correlation matrices were compared with the linear kernels (i.e., similarity matrices) of the environmental covariates (Fig. 1e). Stronger associations between these matrices indicated that the environmental covariate affected the G × E interactions with greater magnitude. These associations were measured using the Pearson’s moment product correlation coefficient (*r*) between the off-diagonal elements of the matrices. Fig. 2 shows the distributions of the −log10 *P* values of *r* calculated for 10 meteorological factors (the complete results are presented in Fig. S4). Generally, YI and SW, and DTT, DTM and SL shared similar *P* values patterns, whereas PR exhibited unique patterns. A total of 555 environmental covariates were significantly detected for the six traits (*P* < 0.05 after Bonferroni correction, Table S17).

**Fig. 2.**
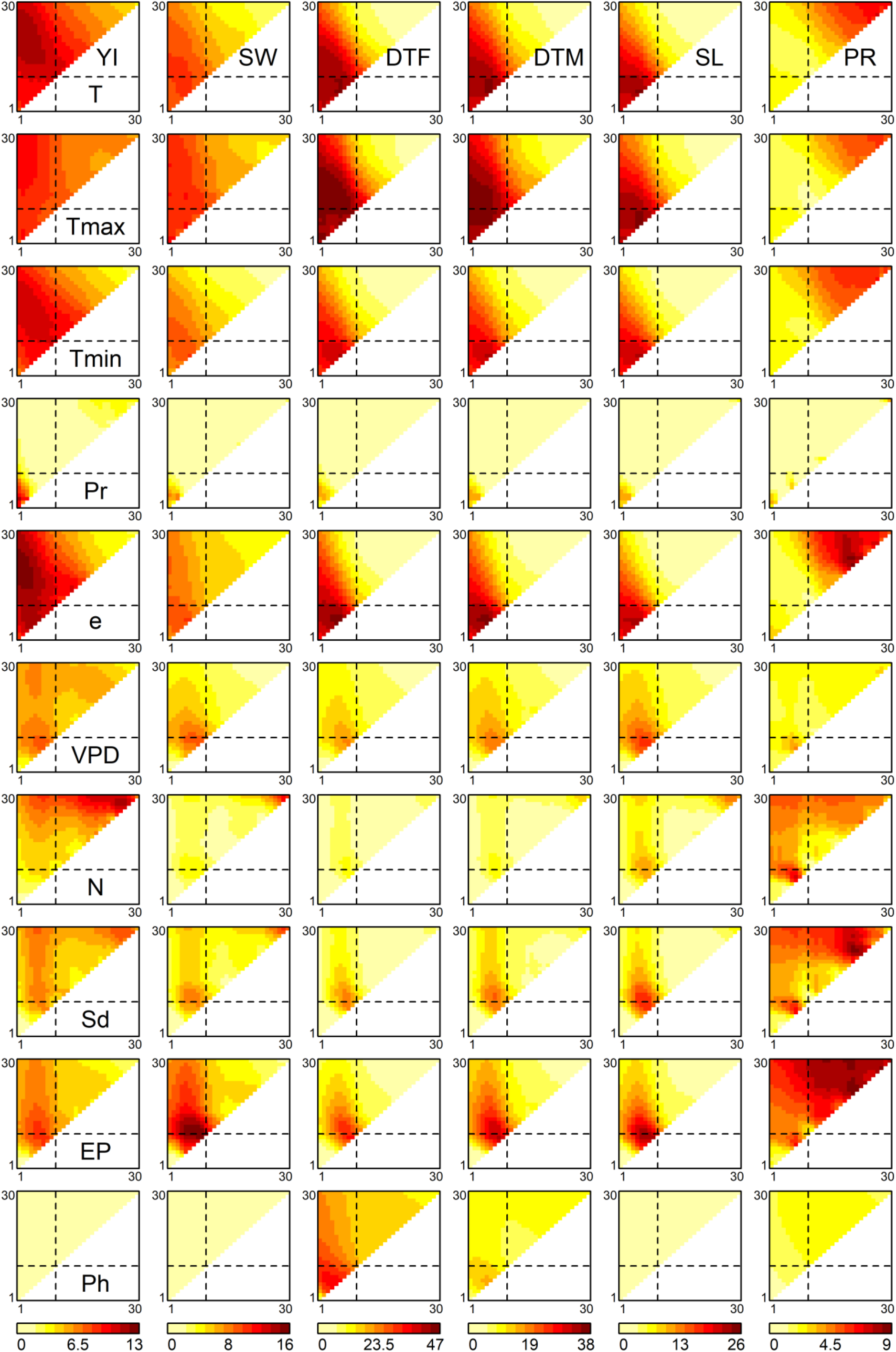
Associations of environmental covariates with genetic correlations between environments. The heat maps represent the −log10 *P* values of correlation coefficients of off-diagonal elements between the similarity matrix of each environmental covariate and the genetic correlation matrix. The diagonal elements of the triangles correspond with the 1^st^ to 30th growth stages, from the lower left to the upper right. The off-diagonal elements correspond to the growth periods that span multiple stages, where the x and y axes denote the start and end of the periods, respectively. The broken lines indicate flowering time. Abbreviations: YI, yield; SW, seed weight; DTF, days to flowering; DTM, days to maturity; SL, stem length; PR, protein content; T, mean temperature; Tmax, maximum temperature; Tmin, minimum temperature; Pr, precipitation; e, vapour pressure; VPD, vapour pressure deficit; N, hours of sunshine; Sd, solar radiation; EP, potential evapotranspiration; Ph, photoperiod.

For YI, three regions in Fig. 2 showed high −log10 *P* values. The first region was detected around the 2^nd^ to 22^nd^ growth stages (upper-left part of the triangles in Fig. 2) of the temperature-related environmental covariates, such as mean temperature, minimum temperature and vapour pressure. The highest −log10 *P* values of these meteorological factors were observed at the 2^nd^ to 22^nd^ growth stages (*r^2^* = 0.036, −log10*P* = 11.554), the 4^th^ to 17^th^ stages (*r^2^* = 0.033, −log10*P* = 10.677) and the 2^nd^ to 19^th^ stages (*r^2^* = 0.039, −log10*P* = 12.460), respectively. The 22^nd^ growth stage corresponded on average to 30.4–34.2 days before maturity. According to the well-known system used to stage soybean development (Fehr & Caviness, 1977), this period largely corresponds to the R5 soybean growth stage (Egli, 2010). Because the R5 stage is defined as the beginning of seed filling (Fehr & Caviness, 1977), it is likely that the temperature-related covariates affect the G × E interactions on YI by modifying the upper limit of the number of seeds. The second region was a period around sowing in relation to precipitation. The highest −log10 *P* value was 10.888 at the 1^st^ to 3^rd^ stages (*r^2^* = 0.034). This stage corresponds to a period from sowing to, on average, 16.5 days after sowing. Soybeans are vulnerable to waterlogging, and, in particular, waterlogging around germination has a severe impact on YI (Kokubun, 2013). In addition, genetic variations exist in root development in flood conditions (Sakazono *et al*., 2014). Thus, our results are reasonable and clearly suggest the importance of precipitation around germination for the G × E interactions regarding YI. The third region was a period just before maturity in relation to hours of sunshine and solar radiation. The highest −log10 *P* value was 11.682 observed for hours of sunshine at the 26^th^ to 29^th^ stages (*r^2^* = 0.037). During this period, these meteorological factors also showed a high −log10 *P* value for SW; the highest value for SW around this period was 11.101 for hours of sunshine at the 29^th^ to 30^th^ stage (*r^2^* = 0.035).

The highest −log10 *P* value for SW was observed in relation to potential evapotranspiration at the 8^th^ to 10^th^ stages (*r*^2^ = 0.050, −log10*P* = 15.715). Potential evapotranspiration (or potential evaporation) can be regarded as the upper limit of evapotranspiration from a crop field (Kuwagata *et al*., 2011), which can reflect photosynthesis activity. The other meteorological factors detected, i.e. mean temperature and maximum temperature (highest *r*^2^ = 0.032 and 0.036 at the 5^th^ stage; −log10*P* = 10.199 and 11.570, respectively) and vapour pressure deficit (*r*^2^ = 0.029 at the 9^th^ stage, −log10*P* = 9.261619), also suggest the relevance of photosynthesis. The influence of these factors started before flowering, when seed filling has not started. Thus, it is likely that these factors affected SW indirectly via the modification of the number of seeds, because soybean shows compensation effects between SW and seed number.

For DTF, the significant environmental covariates included mean temperature at the 6^th^ stage (*r*^2^ = 0.140, −log10*P* = 44.930), minimum temperature at the 6^th^ stage (*r*^2^ = 0.127, −log10*P* = 40.286), maximum temperature at the 5^th^ to 14^th^ stages (*r*^2^ = 0.146, −log10*P* = 46.602) and vapour pressure at the 6^th^ stage (*r*^2^ = 0.145, −log10*P* = 46.353). Photoperiod before flowering was also a significant covariate (*r*^2^ = 0.106 at the 4^th^ to 5^th^ stages, −log10*P* = 33.286). These results are reasonable considering that temperature and photoperiod are the main determinants of soybean flowering time (Sinclair *et al*., 1991). The results obtained for DTM and SL were close to those obtained for DTF, suggesting that the influence of meteorological factors on the G × E interactions for DTM and SL occurred via those of DTF. In other words, if DTF does not show a G × E interaction, DTM and SL will also not show. Finally, the results obtained for SL are reasonable, because the varieties commonly cultivated in Japan exhibit the determinate stem type, in which the prolongation of the stem ends shortly after flowering begins (Bernard, 1972).

The *P* value patterns of PR were different from those of the other traits. Meteorological factors generally exert their effects after flowering, around the 21^st^ to 27^th^ stages, as observed for mean temperature (highest −log*P* = 6.854 at the 27^th^ stage, *r^2^* = 0.062), maximum temperature (−log*P* = 5.544 at the 27^th^ stage, *r^2^* = 0.049), minimum temperature (−log*P* = 6.207 at the 27^th^ stage, *r^2^* = 0.056), vapor pressure (at the 21^st^ stage, −log10*P* = 8.330, *r^2^* = 0.076), solar radiation (−log*P* = 8.605 at the 21^st^ to 24^th^ stages, *r^2^* = 0.079) and potential evapotranspiration (−log*P* = 8.658 at the 21^st^ to 22^nd^ stages, *r^2^* = 0.079). The 21^st^ to 27^th^ stages occurred on average 38–64.6 days after flowering, when the accumulation of protein is past its rapidest growth period (20–40 days after flowering), but is still ongoing (Gayler & Sykes, 1981). The meteorological factors detected potentially affected the G × E interactions of PR via the photosynthetic activity at this stage, which is essential for nitrogen fixation.

To assess the validity of these detected environmental covariates, the detective power of ECGC was verified with simulations. As illustrated in the Methods, the performance of ECGC is affected by *r^2^* and the number of environment *M*. The receiver operating characteristic (ROC) curves obtained under different combinations of *r^2^* and *M* are shown in Fig. 3. As expected, the performance of ECGC gained as *r^2^* and *M* increased. In Fig. 4a and 4b, observed *r^2^* values for each trait are shown for comparison with Fig. 3. The *r^2^* values of the detected environmental covariates were greater than 0.053 for PR (*M* = 30) and greater than 0.018 for the other traits (*M* = 52) (Fig. 4b). The ROC curves under the corresponding combinations of *r^2^* and *M* in simulation results (0.053 and 30, and 0.018 and 50, respectively) suggest that ECGC detected the environmental covariates with reasonable accuracy under these *r^2^* and *M* values.

**Fig. 3.**
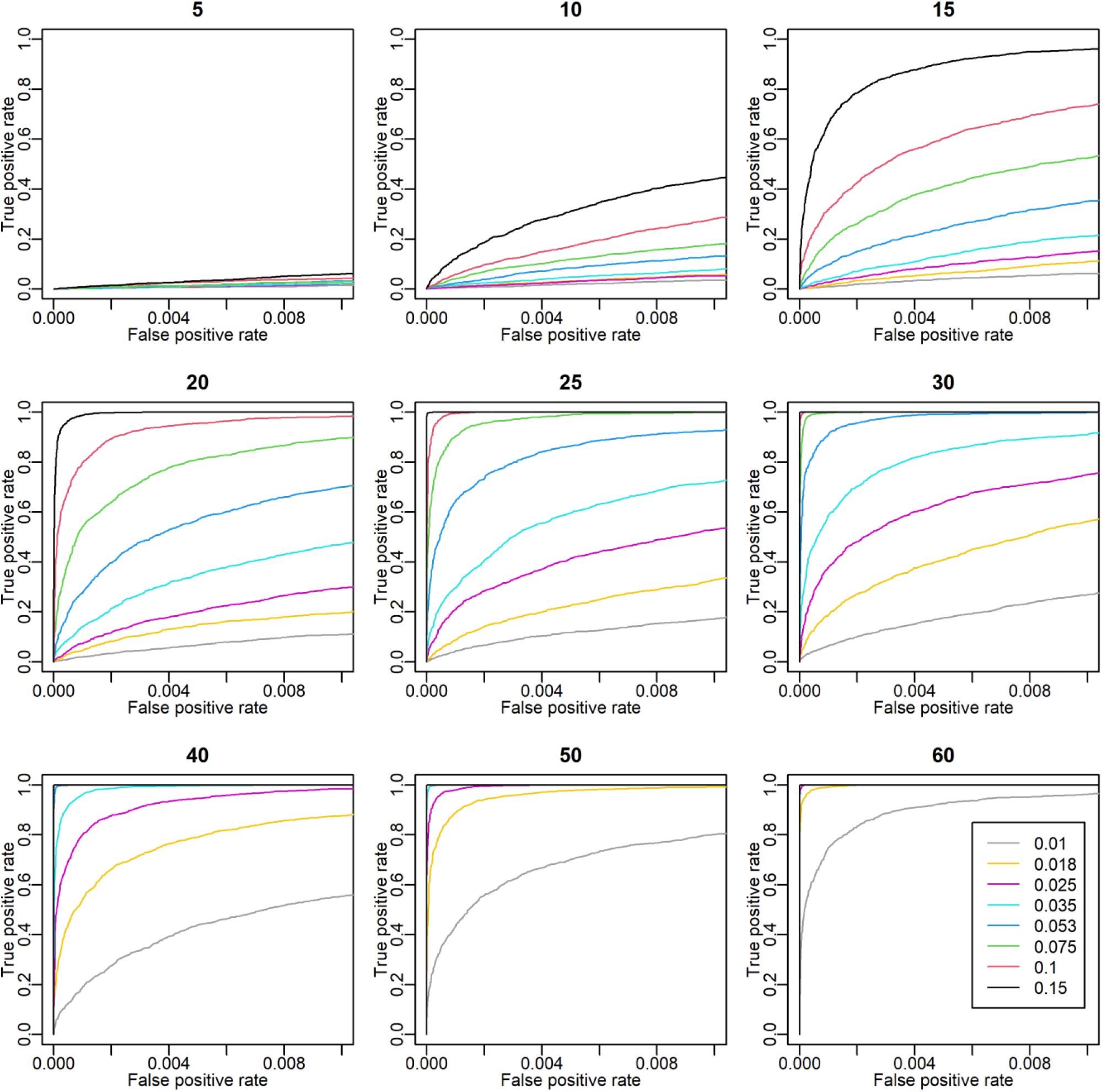
Receiver operating characteristic (ROC) curves in simulation analyses. The titles of the plots denote the number of environments (*M*). The ROC curves of different *r^2^* values are drawn with different colours. The x and y axes are the false positive rate and the true positive rate, respectively. In these simulations, 99 true negatives were simulated for each true positive. Thus, the coordinate (x, y) = (1, 0.01) means that one true positive is detected with 0.99 false positives.

**Fig. 4.**
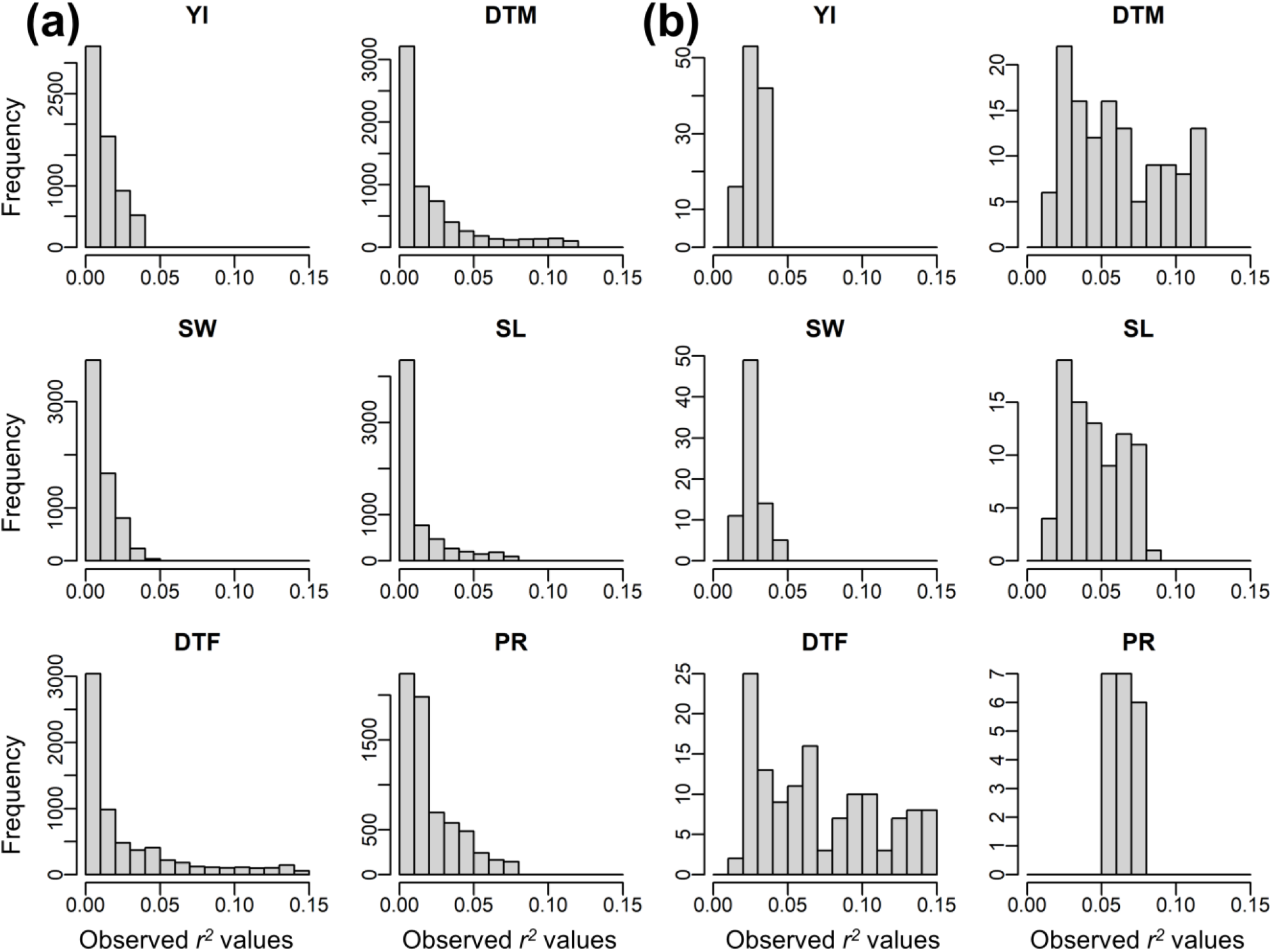
Observed *r^2^* values of off diagonals between the similarity matrix of environmental covariates and the genetic correlation matrix. **(a)** Histograms for all environmental covariates. **(b)** Histograms for the environmental covariates significantly detected. Abbreviations: YI, yield; SW, seed weight; DTF, days to flowering; DTM, days to maturity; SL, stem length; PR, protein content.

In the final step of ECGC, the genetic architecture underpinning the sensitiveness of the varieties to the detected environmental covariates was examined using association mapping (Fig. 1f). For the environmental covariates detected in the fourth step, slopes were estimated for each variety by regressing the additive genetic effects at each environment on the environmental covariates (Table S18). The slope represented the sensitivity of the variety to the changes in the environmental covariates, and genome-wide association (GWA) mapping was conducted on the slopes. By pruning with false discovery rate (FDR) < 0.05 and subsampling analyses, 39 chromosomal regions were significantly detected for traits except for SW (Fig. S5–S10 and Table S19). DTF shared one and two regions with DTM and SL, respectively, and these three traits shared one region (Fig. 5a) which is located at near the known flowering gene (*E2*) (Watanabe *et al*., 2011). On the other hand, YI and PR shared no regions with each other, and nor with DTF, DTM, and SL (Fig. 5a). These results suggest that major genes responsible for the G × E interactions triggered by meteorological factors are generally not pleiotropic. This suggestion can be confirmed by the distributions of the meteorological factor/growth stage combinations where significant associations were detected (Fig. 5b and Fig. S11). For DTF, DTM, and SL, significant associations were generally detected for the temperature-related factors and photoperiod before flowering. For YI, associations were significantly detected for precipitation around sowing, hours of sunshine during maturity, and potential evapotranspiration and vapor pressure deficit (i.e., factors related to photosynthesis) around flowering. For PR, significant associations were mainly detected for temperature-related factors during maturity. That is, the major genes responsible for these traits affect the sensitivities to different meteorological stimuli occurred at different growth stages.

**Fig. 5.**
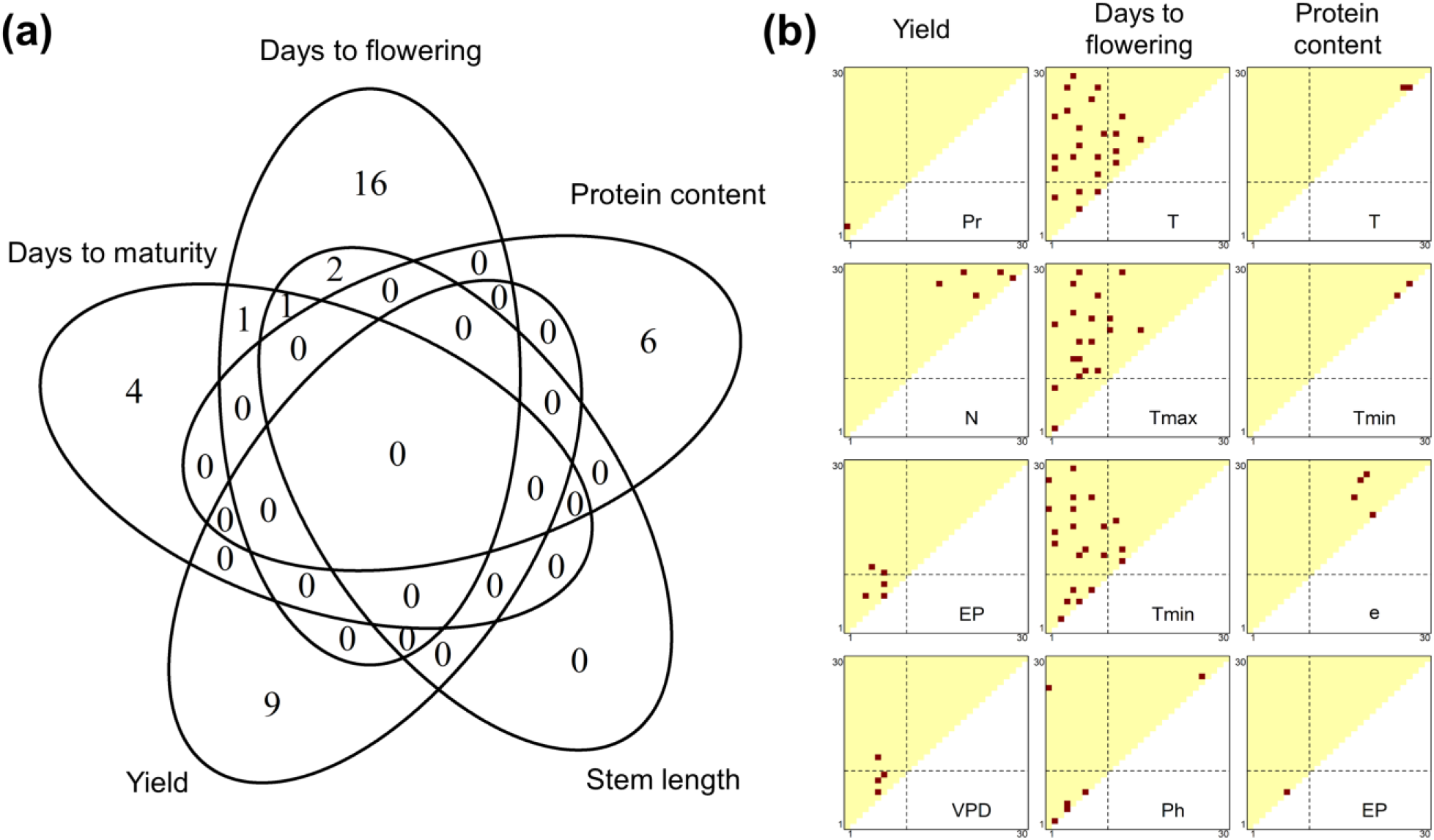
Summary of chromosomal regions detected by genome-wide association mapping. **(a)** Number of chromosomal regions significantly detected by association mapping for each trait. **(b)** Distributions of meteorological factor/growth stage combinations where significant associations were detected. Red dots indicate the combinations with significant associations. See Figure S11 for the other traits and meteorological factors. The diagonal elements of the triangles correspond with the 1^st^ to 30th growth stages, from the lower left to the upper right. The off-diagonal elements correspond to the growth periods that span multiple stages, where the x and y axes denote the start and end of the periods, respectively. The broken lines indicate flowering time. Abbreviations: T, mean temperature; Tmax, maximum temperature; Tmin, minimum temperature; Pr, precipitation; e, vapour pressure; VPD, vapour pressure deficit; N, hours of sunshine; EP, potential evapotranspiration; Ph, photoperiod.

GWA mapping of ECGC could narrow down a genomic region that was suggested to be responsible for the G × E interactions by QTL mapping. Three close regions on chromosome 12 spanning 17.86–18.85 Mbp were detected for the slopes of YI for precipitation at the 1^st^ to 3^rd^ stages (Fig. 6a and 6b, Table S19). These regions include the QTL (14.39–35.1 Mbp) for hypoxia tolerance of Japanese soybean breeds (Van Nguyen *et al*., 2017), which is related to root extension under flood conditions. Gene ontology analyses revealed that two orthologs of *Glyma.12G142900* (18.503–18.505 Mbp) in *Arabidopsis thaliana* (*AT4G27280* and *AT5G54490*) are involved in cell response to hypoxia. In addition, the gene (*Glyma.12G142900*) was reported to show higher expression on root tissues (Phytozome 12, accessed on March 30, 2021), suggesting the gene is a plausible candidate.

**Fig. 6.**
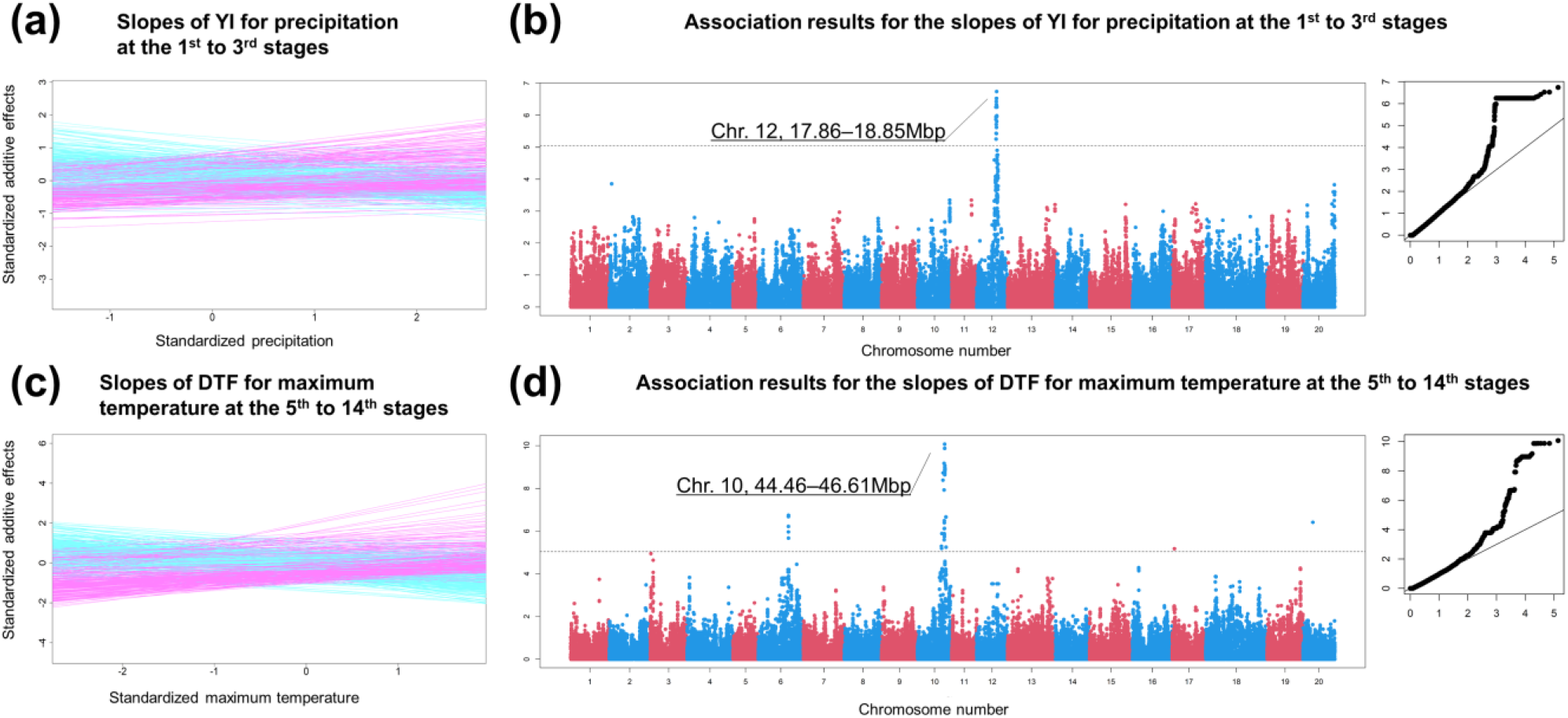
Examples of the genome-wide association mapping results. **(a)** Slopes of YI for precipitation at the 1^st^ to 3^rd^ growth stages. The x and y axes are the environmental covariates and the additive genetic effects, respectively. Both axes are standardised. **(b)** Manhattan and QQ plots of the association analysis of the slopes illustrated in **(a)**. The horizontal dashed line indicates the false discovery rate (0.05) threshold. The x and y axes of the QQ plot are the expected and observed −log10 *P* values, respectively. **(c)** Slopes of days to flowering for maximum temperature at the 5th to 14th growth stages. **(d)** Manhattan and QQ plots for the slopes illustrated in **(c)**.

Strong associations were observed on multiple adjacent regions on chromosome 10 spanning 44.46–46.61 Mbp for various meteorological factors, including temperature-related covariates and photoperiod (Fig. 6c and 6d, Table S19, and Fig. S9). These regions were also detected for DTM and SL, reflecting the similar tendencies of the G × E interactions (Figs. S7 and S10). These regions included *E2* (45.29–45.32 Mbp), which is an analogous gene of *GIGANTEA* of *Arabidopsis* and a major gene for flowering time in soybean (Watanabe *et al*., 2011). These regions also included two orthologs (*Glyma.10G180600* and *Glyma.10G209600*) of *Arabidopsis* flowering genes (*CRY2* and *ELF*6, respectively). Known flowering genes of soybean responsible for photosensitivity, such as *E1* (Xia *et al*., 2012), *E3* (Watanabe *et al*., 2009) and *E4* (Liu *et al*., 2008), were not detected for any of the environmental covariates. It is notable that association mapping using slopes can detect genes with effects that vary according to environmental covariates. Because the loss-of-function alleles of *E1* (Tsubokura *et al*., 2014) were strictly used at higher latitudes (Supplementary Methods 4 and Table S20), the effects at lower latitudes could not be estimated. Thus, these effects would not be reflected in the slopes. Conversely, *E3* and *E4* were probably not involved in the G × E interactions because of constant gene effects across environments. Additional analyses that examined the allele effects of these flowering genes using sommer (Covarrubias-Pazaran, 2016) showed that the *E2* gene effect was the most variable across environments (Supplementary Methods 5 and Fig. S12).

The slopes provide insights on how the G × E interactions were involved in the selection of modern varieties. For example, for YI, varieties with positive slopes for an environmental covariate are expected to be more preferred at environments with greater environmental covariates. In other words, if a trait is directionally preferred, it is expected that the environmental covariates will be correlated with the averages of slopes of the varieties evaluated at each environment. In fact, the correlations were positive for YI and SW (Fig. 7a–c) and often significant (*P* < 0.05 after Bonferroni correction, Table S21). The negative correlations found for PR are attributable to the trade-off between YI and PR (i.e. a higher yield tends to lower PR). These results clearly suggest that varieties that exhibit high YI under the detected environmental covariates have been selected by breeders, intendedly or unintendedly. Conversely, for DTF, DTM and SL, the correlations were more moderate, and no significant correlation was found. This result is reasonable because these traits were not directionally preferred. Rather, a characteristic U-shaped trend was found for temperature-related environmental covariates and photoperiod (Fig. 7d). For DTF, this tendency indicates that a longer DTF is acceptable in lower latitudes (more temperate climates), whereas a shorter DTF is preferred in higher latitudes (less temperate climates). These results are coherent because early flowering is required in cold climates to avoid frost damage.

**Fig. 7.**
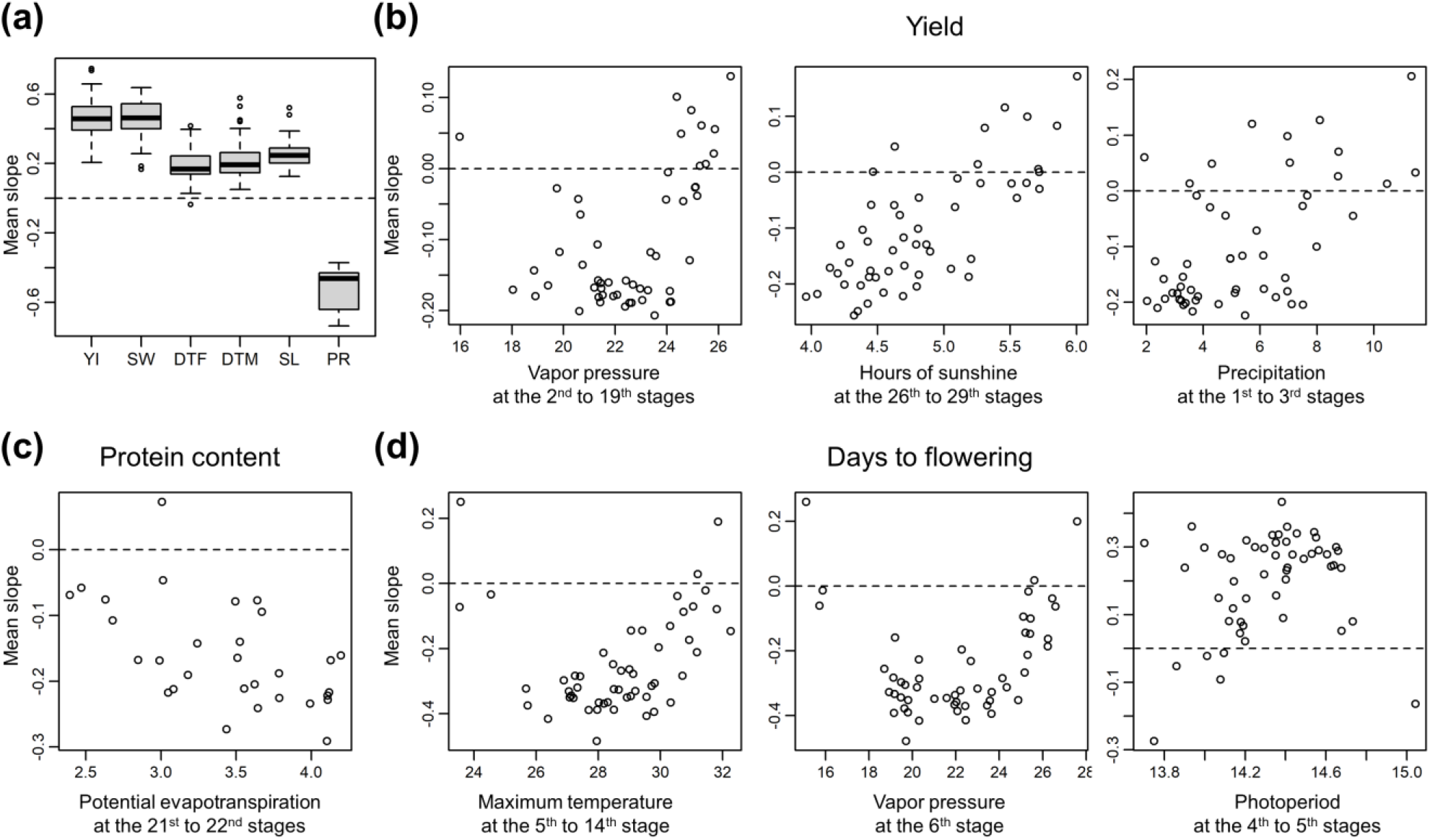
Correlations between environmental covariates and the averages of the slopes of the varieties evaluated at each environment. Positive/negative correlations indicate directional selection for the environmental covariates. **(a)** Weighted averages of slopes for each trait. YI, yield; SW, seed weight; DTF, days to flowering; DTM, days to maturity; SL, stem length; PR, protein content. **(b)** Examples of correlations between the environmental covariates and the averages of the slopes observed for yield. The environment covariates that showed strong associations with the G × E interactions are shown. The red lines are the linear regression lines. **(c)** Example of correlations between the environmental covariates and the averages of the slopes observed for protein content. The environment covariates that showed the highest correlation with the G × E interactions are shown. **(d)** Examples of correlations between the environmental covariates and the averages of the slopes observed for days to flowering. The two environment covariates that exhibited the highest associations with the G × E interactions and photoperiod at the 4th to 5th stages are shown. The red lines were drawn using the local polynomial regression provided by the R package KernSmooth (ver. 2.23-17) (Wand & Jones, 1995).

## Discussion

The base model of ECGC is a conventional model for reaction norm where the genotypic value is divided into a component representing sensitivity to environmental covariates and a component free from them (van Eeuwijk *et al*., 2005; Hayes *et al*., 2016). The novelty of ECGC is to associate similarity matrices of environmental covariates with the genetic correlation matrix, and the idea of ECGC can be derived by extending the base model as described in Materials and Methods. This approach brings the following advantages. First, the environmental covariate search by ECGC is directly related to the G × E interactions whereas the search based on the variance of slopes (sensitivities of genotypes) is not necessarily related to the G × E interactions. Second, because of the properties of mixed models used for estimating genetic correlations between environments, ECGC is applicable to data with missing values and/or unbalanced structure. Genetic correlations between environments can be estimated using mixed models and genomic relationship matrices even when varieties are not overlapped between environments. Estimation of genetic correlation without overlapped records can be found in, for example, estimation of between-sex genetic correlation (Crews & Kemp, 2001). Lastly, genetic covariances/correlations between multiple environments can be estimated in pairwise manner (Fieuws & Verbeke, 2006) and parallelized, which enables ECGC to be scalable with the number of environments. The last two advantages are particularly useful in analysis of large-scale multi-environment data sets and/or historical data sets.

Owing to these advantages, our proposed ECGC was applicable to the large-scale multi-environment data set of soybean, and able to depict comprehensive landscapes on how environment stimuli are involved in the G × E interactions for agronomic traits evaluated in real fields. Moreover, candidate QTLs/genes responsible for the interactions were detected. ECGC also provided interesting insights on how the G × E interactions are related to the selection of modern Japanese varieties. Thus, it can be concluded that ECGC will be a promising approach to understand the G × E interactions and to reveal the gene-by-environment stimuli interactions.

## Acknowledgements

The authors thank Naohiro Yamada of the Nagano Vegetable and Ornamental Crops Experiment Station, Kaori Hirata, Akio Kikuchi, Nobuhiko Oki, Yoshitake Takada and Koji Takahashi of National Agricultural and Research Organization (NARO), Shohei Fujita, Satoshi Kobayashi, Fumiko Kousaka, Hideki Kurosaki, Tomoaki Miyoshi and Keiichi Senda of Hokkaido Research Organization for providing historical data and seeds. The seeds were also provided by Genebank of NARO. The authors also appreciate Makito Hajika of NARO and Hiroyoshi Iwata of the University of Tokyo for their assistance in collaborative works and Ryoma Takeshima, Hiroe Yoshida and Kaori Sasaki of NARO for fruitful discussions. The authors thank two private companies Kumogamisya and Seisyou for digitalising historical booklets.

## Declarations

### Funding

This study was supported by Japan Science and Technology Agency PRESTO (grant number 15656049) and a grant from the Ministry of Agriculture, Forestry and Fisheries of Japan [Smart-breeding system for Innovative Agriculture (BAC1001)].

### Competing interests

The authors declare no competing interests.

### Availability of data and material

Supplementary Tables (Tables S1–21), Supplementary Figures (Figures S1–12), Supplementary Methods (Supplementary Method 1–5), and Supplementary Data (Data 1–4) are available at the bioRxiv web site.

### Code availability

R scripts are available upon requests.

### Author contribution

AO collected and organised historical data, conceived the ECGC framework, conducted all statistical analyses, drafted the manuscript and collaborated on DNA extraction and genotyping. DS extracted DNA and collaborated in drafting the manuscript. AK designed the SNP array, extracted DNA and collaborated on genotyping and drafting the manuscript. SN, TY and JY collaborated on drafting the manuscript. SN suggested the importance of historical data and collaborated on drafting the manuscript.

### Ethics approval

Not applicable

### Consent to participate

Not applicable

### Consent to publication

Not applicable

## Short legends of Supporting Information

**Supplementary Methods 1**: Overview of historical data

**Supplementary Methods 2**: Development of mixed models at each environment

**Supplementary Methods 3**: Estimation of slopes of additive genetic effects on environmental covariates

**Supplementary Methods 4**: Inference of the alleles of flowering genes

**Supplementary Methods 5**: Estimation of the allele substitution effects of flowering genes

**Figure S1**: Results of the clustering of environmental covariates

**Figure S2**: Distribution of the P values obtained from the genome-wide association mapping of the slopes

**Figure S3**: Distributions of estimates of genetic correlations between environments

**Figure S4**: Associations of environmental covariates with genetic correlations between environments

**Figure S5**: Results of the genome-wide association mapping for the slopes obtained for seed weight

**Figure S6**: Results of the genome-wide association mapping for the slopes obtained for yield

**Figure S7**: Results of the genome-wide association mapping for the slopes obtained for stem length

**Figure S8**: Results of the genome-wide association mapping for the slopes obtained for protein content.

**Figure S9**: Results of the genome-wide association mapping for the slopes obtained for days to flowering.

**Figure S10**: Results of the genome-wide association mapping for the slopes obtained for days to maturity.

**Figure S11**: Distributions of meteorological factor/growth stage combinations where significant associations were detected

**Figure S12**: Allele substitution effects of flowering genes (*E2*, *E3* and *E4*) on DTF

**Table S1**: SNP information

**Table S2**: Summary of extracted phenotype data

**Table S3**: Used varieties

**Table S4**: Narrow-sense SNP heritability

**Table S5**: Environmental covariates for yield

**Table S6**: Environmental covariates for seed weight

**Table S7**: Environmental covariates for days to flowering

**Table S8**: Environmental covariates for days to maturity

**Table S9**: Environmental covariates for stem length

**Table S10**: Environmental covariates for protein content

**Table S11**: Genetic correlation of yield between environments

**Table S12**: Genetic correlation of seed weight between environments

**Table S13**: Genetic correlation of days to flowering between environments

**Table S14**: Genetic correlation of days to maturity between environments

**Table S15**: Genetic correlation of stem length between environments

**Table S16**: Genetic correlation of protein content between environments

**Table S17**: Environmental covariates that were significantly correlated with genetic correlations

**Table S18**: Estimated slopes for each variety

**Table S19**: Detected chromosomal regions in genome-wide association mapping and subsampling analysis results

**Table S20**: Unweighted frequencies of flowering gene alleles at each field

**Table S21**: Correlations between environmental covariates at each environment and mean slopes of varieties evaluated at the environments

**Supplementary Data 1**: Phenotypic records and meteorological information

**Supplementary Data 2**: Genomic relationship matrix

**Supplementary Data 3**: Phenotypic records matched between environments for estimating genetic correlations

**Supplementary Data 4**: Matched record IDs

## Notes

### Competing Interest Statement

The authors have declared no competing interest.

