## Supplementary Methods for "A method for identifying environmental stimuli and genes responsible for genotype-by-environment interactions from a large-scale multi-environment data set"

**Contents**

**Methods 1**: Overview of historical data

**Methods 2**: Development of mixed models at each environment

**Methods 3**: Estimation of slopes of additive genetic effects on environmental covariates

**Methods 4**: Inference of the alleles of flowering genes

**Methods 5**: Estimation of the allele substitution effects of flowering genes

### Methods 1. Overview of historical data

The earliest records of modern breeding of soybean in Japan date back to the 1930s, as far as we could investigate. In this study, experimental results from 1961 were subjected to analyses because most weather stations of the Japan Meteorological Agency started observations of major meteorological factors in the 1960s. At each breeding centre, experimental results were released annually as booklets until the 2000s; subsequently, the booklet reports were replaced with digital files, including the Excel and Ichitaro formats, the latter of which is a Japanese application for text editing. The booklets were collected from four breeding centres, i.e. the Central Agricultural Experiment Station of Hokkaido Research Organization, Tohoku Agricultural Research Center of National Agricultural and Research Organization (NARO), Western Region Agricultural Research Center of NARO and Nagano Vegetable and Ornamental Crops Experiment Station. Excel and Ichitaro files were collected from these four centres and the Institute of Crop Science of NARO. The booklets were digitalised and converted to Excel files by commercial services during 2015 to 2018. Subsequently, the digital files, which included experimental results and management conditions (sowing dates, plant density and amount of nitrogen, phosphorus and potassium), were integrated as Excel tables. For correspondence with genomic information, experiments for later generations, typically $F_6$ and later, of which seeds were often available, were subjected to digitalisation and integration. Duplicated records were removed by assessing the years of evaluation, variety names, field names and phenotypic values of yield and stem length. The historical data included 72,829 records of 6,106 varieties evaluated at 440 fields over 55 years (from 1961 to 2015).

Note that each record was an average of the evaluation of multiple plants, with typically two or three replications. However, in statistical analyses, these records were treated as measurements on single plants because the number of replications and plants were often unknown, and varied among fields and years if they were known. It is also notable that, throughout this study, these records were often treated as measurements on single plants for ease of explanations (e.g. Figure 1).

### Methods 2. Development of mixed models at each environment

#### 2.1. Overview

Single-trait mixed models were fitted to the records from each environment. The purposes were (1) to remove variations due to years and management conditions and to adjust the phenotypic values with them, and (2) to estimate the additive genetic effects at the environment (see Table S2 for the data summary at each environment). Here, plant densities and amount of fertiliser are presented as x_y_z, where, for plant density, x, y and z indicate the interval between rows (cm), the interval between strains in a row (cm) and the number of plants per strain; and for fertiliser, x, y and z indicate the amount of nitrogen, phosphate and potassium (kg/10a), respectively. The amount of fertiliser was sometimes unclear because the amount of additional fertiliser was not always described explicitly; in these cases, the amount was regarded as missing (NA). The sowing dates are presented as the days from April 1st.

The models for each environment were developed as follows. The base models included year effects as fixed effects and additive genetic effects as random effects, as follows:

$$\mathbf{Y} = \mathbf{XB} + \mathbf{ZU} + \mathbf{E} \quad (1),$$

where **Y** is a vector of phenotypic values, **X** is an incidence matrix of year effects, **B** is the fixed effect of years, **Z** is a design matrix, **U** is the additive genetic effect and **E** is a vector of residual errors. The covariance structure of **U** was defined using the genome-wide SNPs[1]. When the conditions of a management method (plant density, fertiliser and sowing date) could be regarded as categorical variables, they were modelled as fixed effects, as follows:

$$\mathbf{Y} = \mathbf{XB} + \mathbf{MA} + \mathbf{ZU} + \mathbf{E} \quad (2A),$$

where **M** is an incidence matrix of the effects of the management conditions and **A** is the fixed effect. When the variables could be regarded as continuous, the management conditions were modelled using basis functions, as follows:

$$\mathbf{Y} = \mathbf{XB} + \mathbf{FW} + \mathbf{ZU} + \mathbf{E} \quad (2B),$$

where **F** is a matrix of value output from the basic functions and **W** is a vector of weight of the functions. A row vector of **F** is given as,

$$\mathbf{F}_i = \{f_1(x_i), \quad \cdots, \quad f_n(x_i)\},$$

where $f_j$ is the $j^{\text{th}}$ basis function, $x_i$ is the input value of row $i$ (e.g. the sowing date of variety $i$) and $n$ is the number of functions.

The genotype by management (G × M) interactions were also modelled. When the management conditions were categorical, the model was:

$$\mathbf{Y} = \mathbf{XB} + \mathbf{MA} + \mathbf{ZU} + \mathbf{HV} + \mathbf{E} \quad \text{(3A)},$$

where **H** is a design matrix and **V** is a vector of random effects. **V** was assumed to follow:

$$\mathbf{V} \sim \text{Multivariate Normal}\left(\mathbf{0}, \quad \mathbf{I}_m \otimes \mathbf{G}\sigma_V^2\right),$$

where $\mathbf{I}_m$ is the identity matrix of size $m$, which is the number of categories, **G** is the genetic relationship matrix and $\sigma_V^2$ is the variance component. When the management conditions were continuous, the model was:

$$\mathbf{Y} = \mathbf{XB} + \mathbf{FW} + \mathbf{ZU} + \sum_{j=1}^{n} \mathbf{F}_j \cdot \boldsymbol{Z}\mathbf{Q}_j + \mathbf{E} \quad \text{(3B)},$$

where $\mathbf{F}_j$ is the $j^{\text{th}}$ column vector of **F**, the dot (·) indicates the element-wise product and $\mathbf{Q}_j$ is a vector of random effects for the $j^{\text{th}}$ basis function. $\mathbf{Q}_j$ was assumed to follow:

$$\mathbf{Q}_j \sim \text{Multivariate Normal}\left(\mathbf{0}, \quad \mathbf{G}\sigma_j^2\right).$$

For each management method (plant density, fertiliser and sowing date), models (2) and (3) were compared using AICs, and it was decided whether the G × M interactions were included. Subsequently, using the chosen modelling scheme for each management method (i.e. with or without the interactions), the best combination of methods (e.g. density + fertiliser or fertiliser + sowing) were searched using AIC. Parameter estimation and AIC calculation were conducted using airemlf90[2]. The remainder of this section describes this model-selection process for each environment. The abbreviations of traits are SW, seed weight (g/100 seeds); SL, stem length (cm); YI, yield (kg/a); DTF, days to flowering (days); DTM, days to maturity (days); and PR, protein content (%).

**2.2. F01**

The plant density, fertiliser and sowing date consisted of 14 conditions, five conditions and 36 dates ranging from 50 to 112, respectively. The plant density was transformed to the density per unit area, and then modelled using quadratic curves, as follows:

$$\mathbf{F}_i = \{x_i, \quad x_i^2\}.$$

The effect of the fertiliser was absorbed into the year effect because some conditions were indistinguishable from the year effects. The sowing dates were modelled using cubic splines[3]. The number of splines and knots were 8 and 12, respectively. Thus,

$$\mathbf{F}_i = \{f_1(x_i), \quad \cdots, \quad f_8(x_i)\},$$

where $f$ indicates the spline function. The first and last four knots were arranged to the earliest and latest sowing dates, respectively, to constrain the span of splines[3]. The remaining knots were arranged between them, at equal intervals. This method was common to all the environments for which the sowing dates were modelled using splines.

The AIC values obtained for each management method and trait are presented in Table Sp1. For all traits, models that included the G × M interactions were worse than those without the interactions. Moreover, all models with the interactions failed to converge within the default number of iterations of airemlf90 (5,000). As described below, the models with the interactions often suffered from convergence problems, probably because of an insufficient sample size to support the complex models. For F01, therefore, models without the interactions were selected for all traits and both management methods. The inclusion of the plant density and sowing dates without the G × M interactions yielded the lowest AICs in each trait (Table Sp2). Thus, the model including the density and sowing date effects, but excluding the G × M interactions, was selected.

**Table Sp1** AIC values for each management condition at F01

| | Density | | Sowing date | |
|---|---|---|---|---|
| Trait | Model 2B | Model 3B | Model 2B | Model 3B |
| SW | 3042.1830 | 3068.7756* | 3000.2072 | 3217.8252* |
| SL | 5052.5179 | 5102.8195* | 4983.3719 | 5238.3390* |
| YI | 3563.9963 | 3630.1720* | 3515.1296 | 3782.8403* |
| DTF | 4251.7726 | 4320.7420* | 4098.6910 | 4376.3100* |
| DTM | 4349.3560 | 4400.4659* | 4155.3813 | 4403.9733* |

The lowest values are underlined for each management method.

*Models did not converge.

**Table Sp2** AIC values obtained from models with different management methods at F01

| Trait | Model 1 | + D | + S | + D + S |
|---|---|---|---|---|
| SW | 3152.0639 | 3042.1830 | 3000.2072 | 2994.9552 |
| SL | 5452.7062 | 5052.5179 | 4983.3719 | 4966.1295 |

| | | | | |
|---|---|---|---|---|
| YI | 4035.7022 | 3563.9963 | 3515.1296 | 2102.9812 |
| DTF | 5170.3117 | 4251.7726 | 4098.6910 | 3510.0138 |
| DTM | 5428.2054 | 4349.3560 | 4155.3813 | 4149.2201 |

D and S indicate the density and sowing date, respectively.

The lowest (best) values are underlined.

### 2.3. F02

The plant density, fertiliser and sowing date consisted of three conditions, five conditions and 13 dates ranging from 40 to 65, respectively. The plant density and fertiliser were modelled as fixed effects. The sowing dates were modelled using quadratic curves, and not using splines, because the range was narrow.

The AIC values obtained for each management method are presented in Table Sp3. For all management methods, models including the G × M interactions were worse than those without interactions. Thus, the G × M interactions were not considered. Subsequently, the combinations of management methods were searched using AICs (Table Sp4). As a result, the model that included all management methods was chosen.

**Table Sp3** AIC values for each management condition at F02

| | Density | | Fertiliser | | Sowing date | |
|---|---|---|---|---|---|---|
| Trait | Model 2A | Model 3A | Model 2A | Model 3A | Model 2B | Model 3B |
| SW | 2021.0167 | 2054.6636 | 2012.7094 | 2063.5521 | 2021.2942 | 2043.0269 |
| SL | 2687.1274 | 2718.6261 | 2687.1408 | 2735.2311 | 2695.2464 | 2716.5509 |
| YI | 2115.2979 | 2148.562 | 2106.1844 | 2155.5818 | 2107.8679 | 2129.6604 |
| DTF | 2345.5581 | 2377.5298 | 2339.2051 | 2379.3532 | 2333.1042 | 2350.7157 |
| DTM | 2499.1254 | 2532.2381 | 2483.5045 | 2530.1802 | 2480.8438 | 2498.1562 |

The lowest values are underlined for each management method.

**Table Sp4** AIC values obtained from models with different management methods at F02

| Trait | Model 1 | + D | + F | + S | + D + F | + D + S | + F + S | + D + F + S |
|---|---|---|---|---|---|---|---|---|
| SW | 2025.2599 | 2021.0167 | 2063.5521 | 2043.0269 | 2008.3421 | 2018.2078 | 2007.3502 | 2003.2439 |
| SL | 2699.7367 | 2687.1274 | 2735.2311 | 2716.5509 | 2673.1472 | 2682.9632 | 2682.1078 | 2668.2176 |
| YI | 2123.0094 | 2115.2979 | 2155.5818 | 2129.6604 | 2100.4820 | 2102.9812 | 2093.3760 | 2087.3072 |
| DTF | 2359.9985 | 2345.5581 | 2379.3532 | 2350.7157 | 2325.6281 | 2326.3590 | 2314.2459 | 2304.4848 |

| | | | | | | | | |
|---|---|---|---|---|---|---|---|---|
| DTM | 2506.5943 | 2499.1254 | 2530.1802 | 2498.1562 | 2475.4781 | 2471.3059 | 2464.3322 | 2453.1091 |

D, F and S indicate density, fertiliser and sowing date, respectively.

The lowest (best) values are underlined.

### 2.4. F03

The plant density, fertiliser and sowing date consisted of five conditions, seven conditions and 14 dates ranging from 33 to 85, respectively. The effects of plant density and fertiliser were absorbed into the year effect. The sowing dates were modelled using cubic splines, as described for F01.

AIC values are presented in Table Sp5. For each trait, models including the G × M interactions were better than the models without the interactions. However, as observed in F01, because these models failed to converge within the default number of iterations of airemlf90 (5,000), we avoided the use of the results of the model with the interactions. Instead, the models without the interactions were selected for subsequent analyses, because these models were also better than the base models.

**Table Sp5** AIC values for each management condition at F03

| | | Sowing date | |
|---|---|---|---|
| Trait | Model 1 | Model 2B | Model 3B |
| SW | 1752.0044 | 1676.1747 | 1624.9716* |
| SL | 2816.6115 | 2644.4732 | 2605.3489* |
| YI | 2368.5671 | 2232.7811 | 2187.6439* |
| DTF | 2665.8999 | 1739.6790 | 1663.9495* |
| DTM | 2970.4812 | 2179.3515 | 2113.2196* |

The lowest values are underlined.

*Models did not converge.

### 2.5. F04_D

The data at F04 included two conditions (dense and sparse) as the plant density; thus, the data were divided into two datasets, F04_D and F04_S, which were defined as different environments. The fertiliser and the sowing date of F04_D consisted of two conditions and 14 dates ranging from 68 to 104, respectively. Although the fertiliser consisted of only two conditions, because these two conditions were simultaneously adopted only in 2006, the effects of fertiliser were modelled as the year effects. The sowing dates were modelled using cubic splines, as described for F01. For each trait, models without the G × M interactions were better than those with the interactions and the base models (Table Sp6).

**Table Sp6** AIC values for each management condition at F04_D

| | | Sowing date | |
|---|---|---|---|
| Trait | Model 1 | Model 2B | Model 3B |
| SW | 1480.6452 | 1426.4536 | 1541.7558 |
| SL | 1720.2473 | 1638.8201 | 1752.6496 |
| YI | 1726.0542 | 1628.1234 | 1749.7379 |
| PR | 1237.4889 | 1152.1873 | 1268.4929 |
| DTF | 1730.9428 | 1431.8109 | 1527.4541 |
| DTM | 1866.3623 | 1637.8193 | 1740.4119 |

The lowest (best) values are underlined.

**2.6. F04_S**

The fertiliser and the sowing date of F04_S consisted of two conditions and 14 dates ranging from 75 to 114, respectively. As observed for F04_D, although the fertiliser consisted of only two conditions, because these two conditions were simultaneously adopted only in 2007, the effects of fertiliser were modelled as the year effects. The sowing dates were modelled using cubic splines, as described for F01. For each trait, models without the G × M interactions were the best models (Table Sp7).

**Table Sp7** AIC values for each management condition at F04_D

| | | Sowing date | |
|---|---|---|---|
| Trait | Model 1 | Model 2B | Model 3B |
| SW | 1901.7351 | 1807.397 | 1927.1538 |
| SL | 2922.7259 | 2651.2532 | 2747.4128 |
| YI | 2370.213 | 2246.573 | 2352.105 |
| PR | 1495.2202 | 1388.4733 | 1495.8514 |
| DTF | 2432.961 | 1685.263 | 1722.436 |

| | | | |
|---|---|---|---|
| DTM | 2779.0327 | 2091.5661 | 2206.5631 |

The lowest (best) values are underlined.

**2.7. F14**

The plant density, fertiliser and sowing date consisted of four conditions, eight conditions and 18 dates ranging from 44 to 71, respectively. The plant densities differed in the intervals between the strains. Thus, the intervals were modelled using quadratic curves. The fertiliser was modelled as a fixed effect, whereas the sowing date effects were absorbed into the year effects.

For each management method, models with the G × M interactions were worse than those without the interactions (Table Sp8). Including both the plant density and fertiliser without the G × M interactions yielded the lowest AICs for SL, YI and PR; whereas for SW, DTF and DTM, the inclusion of the fertiliser exclusively, yielded the lowest AICs (Table Sp9). Thus, for the former three traits, the model that included the density and fertiliser effects was selected, whereas for the latter three traits, the model that included the fertiliser effect was selected.

**Table Sp8** AIC values for each management condition at F14

| | Density | | Fertiliser | |
|---|---|---|---|---|
| Trait | Model 2B | Model 3B | Model 2A | Model 3A |
| SW | 8170.371 | 8371.459 | 8154.804 | 8941.904 |
| SL | 11703.23 | 11846.73 | 11816.98 | 12581.74 |
| YI | 9687.95 | 9885.826 | 9930.781 | 10721.1 |
| PR | 3501.383 | 3702.466 | 3490.383 | 4280.7172 |
| DTF | 7835.599 | 8025.853 | 7809.536 | 8586.4951 |
| DTM | 9831.991 | 10030.41 | 9811.265 | 10590.08 |

The lowest values are underlined for each management method.

**Table Sp9** AIC values obtained from models with different management methods at F14

| Trait | Model 1 | + D | + F | + D + F |
|---|---|---|---|---|
| SW | 8168.549 | 8170.371 | 8154.804 | 8157.174 |
| SL | 11909.42 | 11703.23 | 11816.98 | 11654.17 |
| YI | 9982.932 | 9687.95 | 9930.781 | 9664.866 |

| | | | | |
|---|---|---|---|---|
| PR | 3505.383 | 3501.383 | 3490.383 | 3486.383 |
| DTF | 7826.594 | 7835.599 | 7809.536 | 7818.399 |
| DTM | 9829.026 | 9831.991 | 9811.265 | 9815.907 |

D and F indicate the density and fertiliser, respectively.

The lowest (best) values are underlined.

### 2.8. F18

The plant density, fertiliser and sowing date consisted of four conditions, three conditions and 8 dates ranging from 46 to 56, respectively. The plant density was added to the models as a fixed effect, whereas the fertiliser and sowing date were treated as the year effects. The models without the G × M interactions yielded the lowest AIC values for all traits (Table Sp10); thus, they were selected for subsequent analyses.

**Table Sp10** AIC values for each management condition at F18

| | | Density | |
|---|---|---|---|
| Trait | Model 1 | Model 2A | Model 3A |
| SW | 1603.494 | 1590.072 | 1614.178 |
| SL | 2296.655 | 2282.486 | 2301.591 |
| YI | 1753.409 | 1739.89 | 1762.213 |
| DTF | 1895.996 | 1871.665 | 1894.477 |
| DTM | 1989.071 | 1975.000 | 1996.181 |

The lowest values are underlined for each trait.

### 2.9. F19

The plant density and the fertiliser had only one condition. The sowing dates consisted of 22 dates ranging from 26 to 116. The sowing date was modelled using splines, as described for F01. The models with the G × M interactions yielded the lowest AIC values for all traits but SW (Table Sp11). However, the models did not converge, with the exception of that for DTF. Thus, for DTF, the model with the interactions was selected, whereas for the other traits, models without interactions were selected.

**Table Sp11** AIC values for each management condition at F19

| | | Sowing date | |
|---|---|---|---|
| Trait | Model 1 | Model 2B | Model 3B |
| SW | 3311.527 | 3141.978 | 3152.86* |
| SL | 5221.477 | 4862.589 | 4826.331* |
| YI | 4496.984 | 4416.24 | 4411.234* |
| PR | 2509.068 | 2334.44 | 2325.108* |
| DTF | 4417.195 | 3141.678 | 2893.88 |
| DTM | 5767.212 | 4062.131 | 4050.289* |

The lowest values are underlined for each trait.
*Models did not converge.

## 2.10. F21

Both the plant density and the fertiliser consisted of eight conditions. The sowing dates consisted of 31 dates ranging from 64 to 119. The sowing date was modelled using splines, as described for F01. The models with the G × M interactions yielded the lowest AIC values for PR and DTF, whereas the models without the interactions yielded the lowest for the remaining traits (Table Sp12). However, because the models with the interactions did not converge, the models without the interactions were selected for all traits.

**Table Sp12** AIC values for each management condition at F21

| | | Sowing date | |
|---|---|---|---|
| Trait | Model 1 | Model 2B | Model 3B |
| SW | 1467.893 | 1429.301 | 1431.424* |
| SL | 2241.805 | 2186.105 | 2190.856* |
| YI | 1961.558 | 1908.521 | 1912.742* |
| PR | 824.7133 | 778.2164 | 771.5191* |
| DTF | 1809.297 | 1579.229 | 1575.351* |
| DTM | 2181.5907 | 1784.1958 | 1793.8385* |

The lowest values are underlined for each trait.
*Models did not converge.

## 2.11. F27_D

The data at F27 were divided into two sets according to the plant density, i.e. dense (F27_D) and sparse (F27_S). The fertiliser of F27_D consisted of three conditions, which were absorbed into the year effects. The sowing dates, which ranged from 63 to 97, were modelled using cubic splines, as described for F01. For all traits, the models with the sowing date effects but without the G × M interactions yielded the lowest AIC values (Table Sp13).

**Table Sp13** AIC values for each management condition at F27_D

| | | Sowing date | |
|---|---|---|---|
| Trait | Model 1 | Model 2B | Model 3B |
| SW | 2623.711 | 2604.287 | 2907.379 |
| SL | 3005.028 | 2984.849 | 3299.296 |
| YI | 2671.586 | 2650.957 | 2997.626 |
| PR | 2040.373 | 2029.373 | 2373.638 |
| DTF | 2572.863 | 2344.663 | 2652.719 |
| DTM | 2976.736 | 2940.729 | 3253.466 |

The lowest values are underlined for each trait.

**2.12. F27_S**

The fertiliser of F27_S consisted of three conditions that were absorbed into the year effects. The sowing dates, which ranged from 37 to 78, were modelled using cubic splines. For all traits, the models with the sowing date effects but without the G × M interactions yielded the lowest AIC values (Table Sp14).

**Table Sp14** AIC values for each management condition at F27_S

| | | Sowing date | |
|---|---|---|---|
| Trait | Model 1 | Model 2B | Model 3B |
| SW | 4059.107 | 4010.586 | 4316.941 |
| SL | 5737.045 | 5682.37 | 6001.286 |
| YI | 4889.413 | 4824.363 | 5139.919 |
| PR | 2231.696 | 2172.287 | 2528.741 |
| DTF | 4691.164 | 4110.358 | 4423.492 |
| DTM | 5370.723 | 5075.687 | 5380.421 |

The lowest values are underlined for each trait.

**2.13. F29**

The plant density and the fertiliser consisted of four conditions. These conditions were indistinguishable from the year effects; thus, their effects were absorbed into the year effects. The sowing date consisted of 25 dates ranging from 66 to 113, and was modelled using cubic splines, as described for F01. The inclusion of the sowing date effects improved the model (Table Sp15), whereas the inclusion of the G × M interactions only improved the model for DTF. Moreover, because the models with the interactions did not converge for the traits, including DTF, the models without the interactions were selected.

**Table Sp15** AIC values for each management condition at F29

| | | Sowing date | |
|---|---|---|---|
| Trait | Model 1 | Model 2B | Model 3B |
| SW | 985.0245 | <u>949.1639</u> | 966.0412* |
| SL | 1400.413 | <u>1344.698</u> | 1361.121* |
| YI | 1188.658 | <u>1136.669</u> | 1162.296* |
| PR | 498.3626 | <u>464.7469</u> | 478.5859 |
| DTF | 1237.57 | 953.4021 | <u>875.9031*</u> |
| DTM | 1525.291 | <u>1196.959</u> | 1245.283* |

The lowest values are underlined for each trait.

*Models did not converge.

**2.14. F30**

The plant density, fertiliser and sowing date consisted of nine conditions, six conditions and 27 dates ranging from 56 to 95, respectively. The fertiliser effects were absorbed into the year effects because they were indistinguishable. The plant density was modelled as a fixed effect, whereas the sowing date effects were modelled using cubic splines, as described in F01.

For each management method (plant density and sowing date), the models with the G × M interactions were worse than those without the interactions (Table Sp16). The inclusion of both the plant density and the sowing date without the G × M interactions yielded the lowest AICs for all traits (Table Sp17).

**Table Sp16** AIC values for each management condition at F30

| | Density | | Sowing date | |
|---|---|---|---|---|
| Trait | Model 2A | Model 3A | Model 2B | Model 3B |
| SW | 2727.155 | 2918.393 | 2701.481 | 2942.512 |
| SL | 3881.645 | 4071.645 | 3821.31 | 4068.587 |
| YI | 3018.938 | 3209.442 | 2985.041 | 3230.614 |
| DTF | 2884.001 | 3075.165 | 2647.859 | 2891.85 |
| DTM | 3439.574 | 3630.802 | 3259.505 | 3516.088 |

The lowest values are underlined for each management method.

**Table Sp17** AIC values obtained from models with different management methods at F30

| Trait | Model 1 | + D | + S | + D + S |
|---|---|---|---|---|
| SW | 2747.131 | 2727.155 | 2701.481 | 2678.093 |
| SL | 3948.907 | 3881.645 | 3821.31 | 3780.626 |
| YI | 3057.589 | 3018.938 | 2985.041 | 2950.343 |
| DTF | 3063.325 | 2884.001 | 2647.859 | 2604.186 |
| DTM | 3652.693 | 3439.574 | 3259.505 | 3229.986 |

D and S indicate the density and sowing date, respectively.
The lowest (best) values are underlined.

## 2.15. F34_D

The data at F34 were divided into two data sets according to the plant density, i.e. dense (F34_D) and sparse (F34_S). The fertiliser consisted of two conditions, which were indistinguishable from the year effects; thus, their effects were absorbed into the year effects. The sowing date consisted of 18 dates ranging from 48 to 103 and was modelled using cubic splines, as described for F01. The models that included the sowing date effects without the G × M interactions were better than the models with the G × M interactions for all traits (Table Sp18).

**Table Sp18** AIC values for each management condition at F34_D

| | | Sowing date | |
|---|---|---|---|
| Trait | Model 1 | Model 2B | Model 3B |
| SW | 5473.603 | 5367.076 | 5468.295 |
| SL | 7298.3 | 7015.35 | 7113.368 |

| | | | |
|---|---|---|---|
| YI | 5953.777 | 5841.93 | 5877.806 |
| PR | 4060.772 | 3915.081 | 4139.809 |
| DTF | 6379.479 | 5753.834 | 5768.883 |
| DTM | 6928.906 | 6466.64 | 6577.857 |

The lowest values are underlined for each trait.

### 2.16. F34_S

The fertiliser consisted of two conditions and its effects were absorbed into the year effects. The sowing date consisted of 15 dates ranging from 48 to 107 and was modelled using cubic splines. The models that included the sowing date effects without the G × M interactions were better than the models with the G × M interactions for all traits (Table Sp19).

**Table Sp19** AIC values for each management condition at F34_S

| | | Sowing date | |
|---|---|---|---|
| Trait | Model 1 | Model 2B | Model 3B |
| SW | 6434.538 | 6357.721 | 6477.721 |
| SL | 9633.348 | 9352.52 | 9402.386 |
| YI | 7597.851 | 7410.378 | 7512.847 |
| PR | 4386.634 | 4356.416 | 4529.635 |
| DTF | 8046.311 | 7320.69 | 7424.064 |
| DTM | 8807.109 | 8170.769 | 8253.725 |

The lowest values are underlined for each trait

### 2.17. F41

The plant density, fertiliser and sowing date consisted of five conditions, 13 conditions and 19 dates ranging from 32 to 90, respectively. The plant densities per unit area ($m^2$) were calculated and modelled using quadratic curves. The sowing date was modelled using cubic splines, whereas the fertiliser effects were absorbed into the year effects.

The models with the G × M interactions were suggested for SL of the sowing date (Table Sp20). However, because the model did not converge within the default number of iterations of the programme, the model without the interactions was adopted. The inclusion of both the plant density and sowing date without the G × M interactions yielded the lowest AICs for all traits (Table Sp21).

**Table Sp20** AIC values for each management condition at F41

| | Density | | Sowing date | |
|---|---|---|---|---|
| Trait | Model 2B | Model 3B | Model 2B | Model 3B |
| SW | 1682.048 | 1690.672* | 1633.283 | 1673.698* |
| SL | 2542.197 | 2550.047 | 2487.835 | 2483.162* |
| YI | 2042.009 | 2049.307* | 1993.373 | 2028.041* |
| DTF | 2105.78 | 2115.148* | 1898.7 | 1926.5* |
| DTM | 2304.014 | 2312.105 | 2207.663 | 2243.782* |

The lowest values are underlined for each management method

*Model did not converge

**Table Sp21** AIC values obtained from models with different management methods at F41

| Trait | Model 1 | + D | + S | + D + S |
|---|---|---|---|---|
| SW | 1693.965 | 1682.048 | 1633.283 | 1632.721 |
| SL | 2558.78 | 2542.197 | 2487.835 | 2481.938 |
| YI | 2058.089 | 2042.009 | 1993.373 | 1989.102 |
| DTF | 2534.999 | 2105.78 | 1898.7 | 1891.339 |
| DTM | 2715.438 | 2304.014 | 2207.663 | 2203.414 |

D and S indicate the density and sowing date, respectively.

The lowest (best) values are underlined.

#### 2.18. Environments in which the base models were fitted

Table Sp22 presents the environments in which the base models (model 1) were fitted because the conditions of all management methods were indistinguishable from the year effects. The data at F07 included two conditions (early and late) of sowing date, and the data were divided into two data sets, F07_E and F07_L, which were defined as different environments. Similarly, the data at F08 included two conditions (early and late) of sowing date, and the data were divided into two data sets, F08_E and F08_L. The data at F09 included two conditions (dense and sparse) of plant density, which were denoted as F09_D and F09_S, respectively. The evaluation at F22 was performed in two conditions, i.e. dense plant density (75_10_2)/late sowing (81–96) and sparse plant density (75_20_2)/early sowing

(54–62), which were denoted as F22_D and F22_S, respectively. The fertiliser was 2_6_8 in both conditions throughout the study period. The data at F24 were divided into three sets according to the sowing date and the plant density; i.e. early date/dense (F24_ED), early date/sparse (F24_ES) and late date (F24_L).

**Table Sp22** Environments and number of conditions of each management method

| Environment ID | Plant density | Fertiliser | Sowing date (earliest–latest) |
|---|---|---|---|
| F05 | 3 | 5 | 22 (68–114) |
| F06 | 12 | 11 | 22 (27–79) |
| F07_E | 1 | 3 | 14 (33–74) |
| F07_L | 2 | 6 | 10 (84–98) |
| F08_E | 5 | 12 | 11 (49–60) |
| F08_L | 2 | 9 | 8 (78–92) |
| F09_D | 1 | 15 | 17 (53–98) |
| F09_S | 2 | 20 | 11 (49–64) |
| F10 | 20 | 14 | 23 (51–108) |
| F11 | 8 | 9 | 17 (52–73) |
| F12 | 9 | 14 | 19 (43–90) |
| F13 | 8 | 11 | 18 (67–92) |
| F15 | 2 | 23 | 26 (49–106) |
| F16 | 6 | 6 | 14 (76–93) |
| F17 | 3 | 9 | 15 (48–68) |
| F20 | 9 | 18 | 13 (76–110) |
| F22_D | 1 | 1 | 9 (81–96) |
| F22_S | 1 | 1 | 7 (54–62) |
| F23 | 1 | 11 | 9 (50–59) |
| F24_ED | 1 | 3 | 5 (59–63) |
| F24_ES | 1 | 3 | 5 (59–63) |
| F24_L | 1 | 5 | 7 (90–96) |
| F25_ED | 1 | 4 | 6 (55–65) |
| F25_ES | 1 | 5 | 9 (55–65) |
| F25_L | 1 | 4 | 4 (93–96) |
| F26 | 12 | 12 | 18 (75–102) |
| F28 | 14 | 20 | 26 (62–116) |
| F31 | 6 | 4 | 13 (62–89) |
| F32 | 16 | 15 | 20 (60–98) |

| | | | |
|---|---|---|---|
| F33 | 1 | 6 | 10 (76–89) |
| F35 | 1 | 2 | 5 (62–67) |
| F36 | 10 | 3 | 15 (64–80) |
| F37 | 7 | 4 | 17 (50–72) |
| F38 | 5 | 4 | 16 (71–115) |
| F39 | 5 | 3 | 16 (51–81) |
| F40 | 6 | 3 | 8 (55–67) |

**Methods 3. Estimation of slopes of additive genetic effects on environmental covariates**

The estimates of the additive genetic effects of a variety at the 52 environments (30 in the case of PR) were regressed on the environmental covariates and the slopes were estimated using a maximum likelihood approach. This process is written for variety *i* as:

$$\hat{u}_{i,j} = \mu_i + x_j \beta_i + \varepsilon_{i,j},$$

where $\hat{u}_{i,j}$ is the estimate of the additive genetic effect at environment *j* which is scaled with the estimate of the additive genetic standard deviation ($\hat{\sigma}_{u_j}$), $\mu_i$ is the intercept, $x_j$ is the standardized environmental covariate at environment $j$, $\beta_i$ is the slope and $\varepsilon_{i,j}$ is the residual error. The residual error follows a normal distribution,

$$\varepsilon_{i,j} \sim \text{Normal}\left(0,\ \ \sigma^2_{\tilde{u}_j} + \sigma^2_{i,j}\right),$$

where $\sigma^2_{\tilde{u}_j}$ is the variance of the parts of the additive genetic effects that were unexplainable by the environmental covariate (see Methods in the main text) and $\sigma^2_{i,j}$ is the prediction error variance (i.e., uncertainty of additive genetic effects or posterior variance) which is scaled with $\hat{\sigma}_{u_j}$.

In this procedure, for stable computation, two assumptions in ECGC were simplified. First, the slope ($\beta_i$) was treated as a fixed effect and estimated for each variety independently. Second, $\sigma^2_{\tilde{u}_j}$ is assumed to be common to each environment (i.e., $\sigma^2_{\tilde{u}}$). Without these simplifications, estimation of the slopes was unstable because of high dimensional data (624 varieties and 52 environments). Here, $\sigma^2_{i,j}$ is a known value and the remaining values ($\mu_i$, $\beta_i$ and $\sigma^2_{\tilde{u}}$) are parameters to be estimated. The log likelihood is as follows:

$$\mathrm{L} = -\frac{1}{2}\sum_j \log\left(\sigma^2_{\tilde{u}} + \sigma^2_{i,j}\right) - \frac{1}{2}\sum_j \frac{\left(\hat{u}_{i,j} - \mu_i - x_j \beta_i\right)^2}{\sigma^2_{\tilde{u}} + \sigma^2_{i,j}}.$$

The maximum likelihood estimates of these parameters were obtained by iteratively maximising the log likelihood with regard to each parameter. $\beta_i$ was updated as:

$$\hat{\beta}_i = \sum_j \frac{x_j\left(\hat{u}_{i,j} - \hat{\mu}_i\right)}{\sigma^2_{\tilde{u}} + \sigma^2_{i,j}} \Big/ \sum_j \frac{x_j^2}{\sigma^2_{\tilde{u}} + \sigma^2_{i,j}},$$

and $\mu_i$ was updated as:

$$\hat{\mu}_i = \sum_j \frac{\hat{u}_{i,j} - x_j \hat{\beta}_i}{\sigma^2_{\tilde{u}} + \sigma^2_{i,j}} \Big/ \sum_j \frac{1}{\sigma^2_{\tilde{u}} + \sigma^2_{i,j}}.$$

These equations were obtained by setting the first derivatives of the loglikelihood as zero. Because $\sigma^2_{\tilde{u}}$ does not have a simple solution using this approach, $\sigma^2_{\tilde{u}}$ was updated by maximising the log likelihood directly using the Brent algorithm implemented in the R function optim. The initial values were 0 for $\beta_i$ and $\mu_i$ and 1 for $\sigma^2_{\tilde{u}}$. The iteration stopped when the summation of the absolute values of the changes of these parameters was $< 1e{-8}$. Usually, the iteration stopped quickly (<100 iterations).

**Methods 4. Inference of the alleles of flowering genes**

Because the SNP array used here does not include the alleles of known flowering genes (*E1*, *E2*, *E3* and *E4*), we inferred the alleles of these genes for the used varieties from the SNP genotypes. The alleles of the four flowering genes were reported for 63 varieties[1], and 26 varieties were overlapped with our study. Using these 26 varieties as the reference, the correspondence between the haplotypes of SNPs located around the genes and the reported alleles was investigated. Haplotypes were constructed by gradually expanding the target region in both the 5′ and 3′ directions of the genes, such that alleles were distinguished by haplotypes.

For *E1*, 12 haplotypes were constructed from 209 SNPs that were located within the ± 600 kbp region of the gene (*Glyma.06G207800*, 20207077–20207940 bp on chromosome 6). Among the 12 haplotypes found, one corresponded to the *e1-nl* allele, one corresponded to the *e1-as* allele and 10 corresponded to the *E1* allele. Although a variety (V619) carrying the *E1* allele exhibited the haplotype corresponding to the *e1-nl* allele, because this variety could not be distinguished from the other varieties with the *e1-nl* allele even when the span of the haplotypes was expanded further, we assigned the *e1-nl* allele to this variety. Because the 79 varieties included in our study had haplotypes that were not represented in the reference, we could not determine the alleles of these varieties. For *E2*, five haplotypes were constructed from 30 SNPs located within the ± 50 kbp region of the gene (*Glyma.10G221500*, 45294737–45316113 bp on chromosome 10). One, one and three haplotypes corresponded to the *E2-dl*, *E2-in* and *e2-ns* alleles, respectively. The haplotypes of 13 varieties were not represented in the reference and were not determined. For *E3*, three haplotypes were constructed from four SNPs located within the ± 10 kbp region of the gene (*Glyma.19G224200*, 47633059–47641958 bp on chromosome 19). These haplotypes corresponded to the *E3-Mi*, *E3-Ha* and *e3-tr* alleles, respectively. Because a variety carrying the *e3-tr* allele (V163) exhibited the haplotype of *E3-Mi*, we assigned the *E3-Mi* allele to this variety. The haplotypes of five varieties were not represented in the reference. For *E4*, 10 haplotypes were constructed from 18 SNPs located within the ± 300 kbp region of the gene (*Glyma.20G090000*, 33236018–33241692 bp on chromosome 20). One haplotype corresponded to the *e4-SORE-I* allele, and the remaining haplotypes corresponded to the *E4* allele. The haplotypes of 220 varieties were not represented in the reference.

### Methods 5. Estimation of the allele substitution effects of flowering genes

The allele substitution effects of flowering genes (*E2*, *E3* and *E4*) were estimated using mixed models at each environment. *E1* was omitted from this analysis because the gene was fixed with the functional allele (*E1*) at most fields (Table S18). The alleles of *E2*, *E3* and *E4* were inferred as described in the previous section. The mixed models can be written as:

$$\tilde{\mathbf{Y}} = \mathbf{XB} + \mathbf{ZU} + \mathbf{E},$$

where $\tilde{\mathbf{Y}}$ is a vector of phenotypic values adjusted for year and management conditions (as described in Section 1), **X** is an incidence matrix representing flowering gene alleles, **B** is the fixed effects of alleles, **Z** is a design matrix, **U** is the additive genetic effect and **E** is a vector of residual errors. The covariance structure of **U** was defined using the genome-wide SNPs[1]. Because the loss-of-function alleles (*e2-ns*, *e3-tr* and *e4-SORE-I*) are responsible for early flowering, we estimated the effects of these alleles against functional alleles, *E2-dl* and *E2-in*, *E3-Mi* and *E3-Ha* and *E4*, respectively. Thus, **X** has four columns that represent the intercept, *e2-ns*, *e3-tr* and *e4-SORE-I*, respectively. The parameters were estimated using AI-REML implemented in the R package sommer[1].
