## Supplementary Figures for "A method for identifying environmental stimuli and genes responsible for genotype-by-environment interactions from a large-scale multi-environment data set"

Akio Onogi*[1], Daisuke Sekine[2], Akito Kaga[3], Satoshi Nakano[4], Tetsuya Yamada[3], Jianming Yu[5], Seishi Ninomiya[6]

[1] Department of Plant Life Science, Faculty of Agriculture, Ryukoku University, Otsu, Shiga, 520-2194, Japan
[2]Institute of Vegetable and Floriculture Science, National Agriculture and Food Research Organization, Tsu, Mie 514-2392, Japan
[3]Institute of Crop Science, National Agriculture and Food Research Organization, Tsukuba, Ibaraki 305-8518, Japan
[4]Institute for Agro-Environmental Sciences, National Agriculture and Food Research Organization, Tsukuba, Ibaraki 305-8604, Japan
[5]Department of Agronomy, Iowa State University, Ames, IA 50011, USA
[6]Graduate School of Agricultural and Life Science, The University of Tokyo, Nishitokyo, Tokyo 188-0002, Japan

**Corresponding author:**
Akio Onogi
Postal address: 1-5, Yokotani, Oe-cho, Seta, Otsu, Shiga, 520-2194, Japan


### Abbreviations

#### Trait

| | |
|---|---|
| **YI** | yield |
| **SW** | seed weight |
| **DTF** | days to flowering |
| **DTM** | days to maturity |
| **SL** | stem length |
| **PR** | protein content; |

#### Meteorological factor

| | | | |
|---|---|---|---|
| **T** | mean temperature | **u** | wind speed |
| **Tmax** | maximum temperature | **u10max** | maximum wind speed |
| **Tmin** | minimum temperature | **N** | hours of sunshine |
| **Pr** | precipitation | **Ss** | solar radiation |
| **e** | vapour pressure | **EP** | potential evapotranspiration |
| **VPD** | vapour pressure deficit | **Ph** | photoperiod. |
| **RH** | relative humidity | | |
| **RHmin** | minimum relative humidity | | |

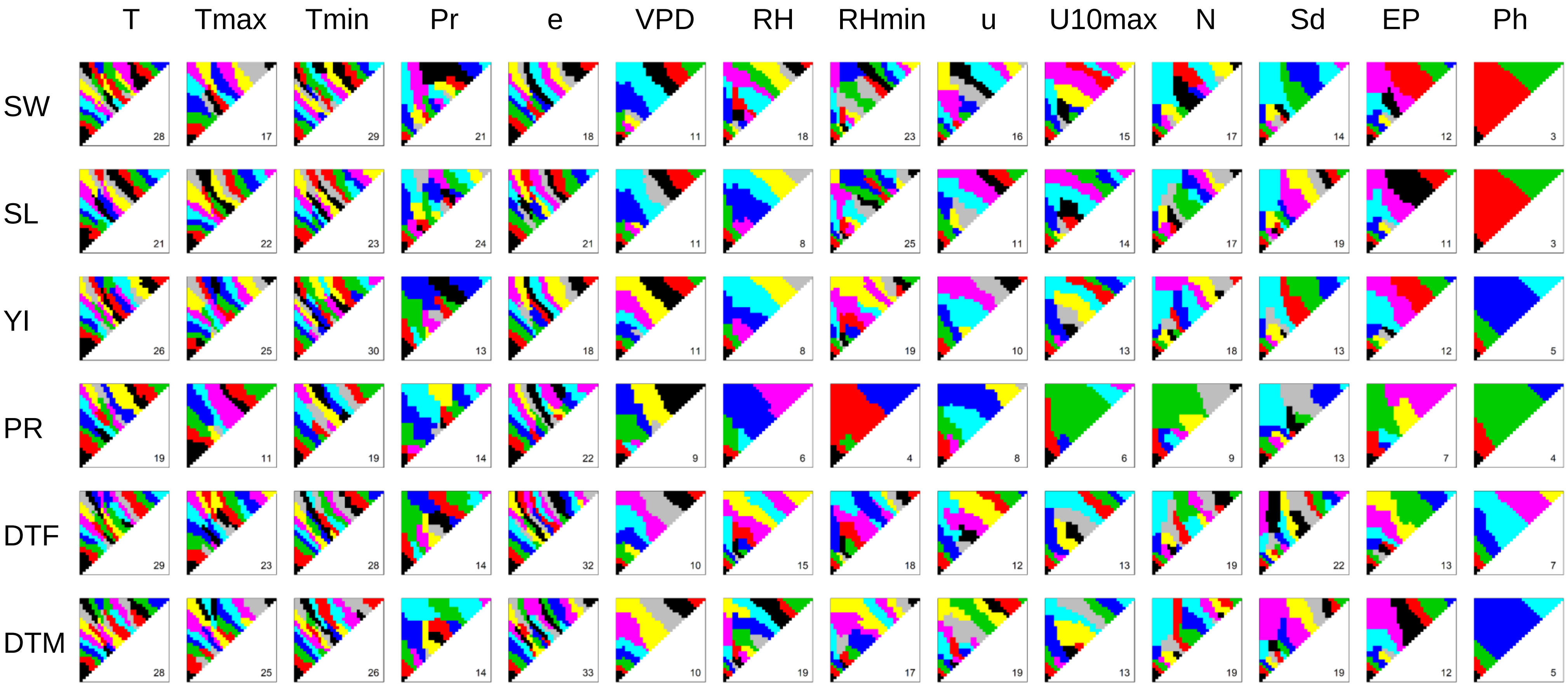


**Figure S1 Results of the clustering of environmental covariates.** The diagonal boxes of the triangles correspond to the 1st to 30th growth stages, from the lower left to the upper right. The off-diagonal elements correspond to the growth periods that span multiple stages, where the x and y axes denote the start and end of the periods, respectively. Note that, although neighbouring clusters are coloured using different colours, the same colour does not indicate the same cluster. The number of clusters is shown in the plots.

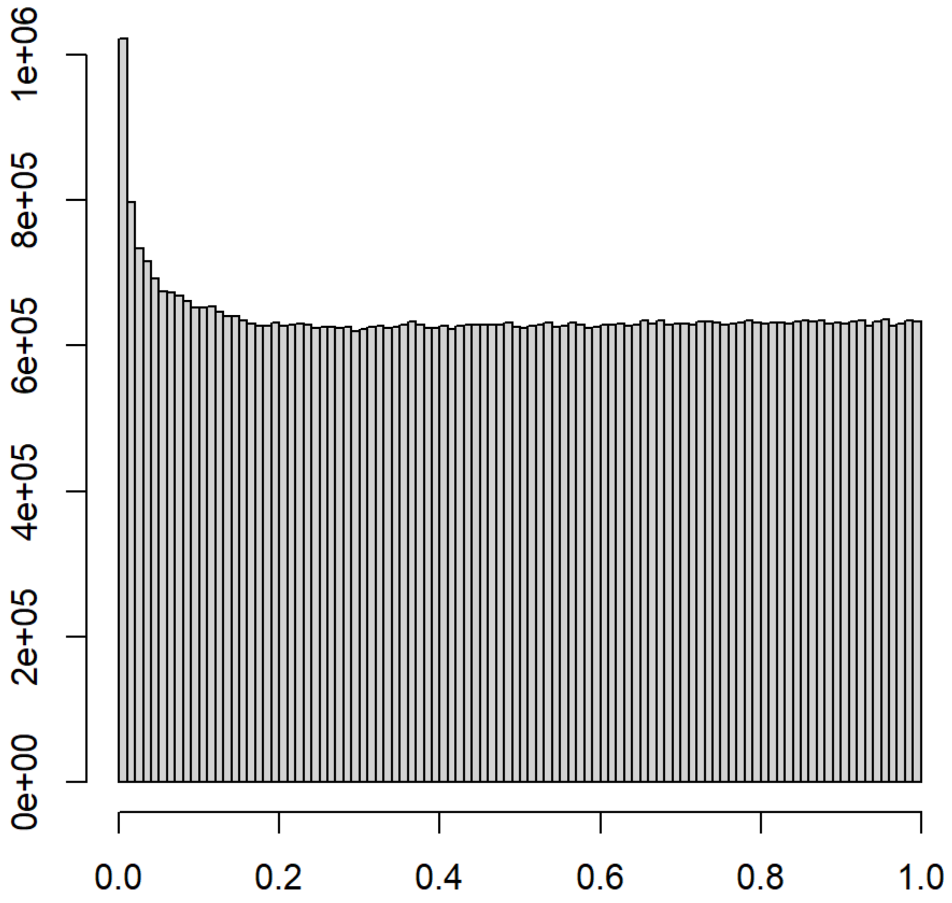


**Figure S2 Distribution of the *P* values obtained from the genome-wide association mapping of the slopes.**

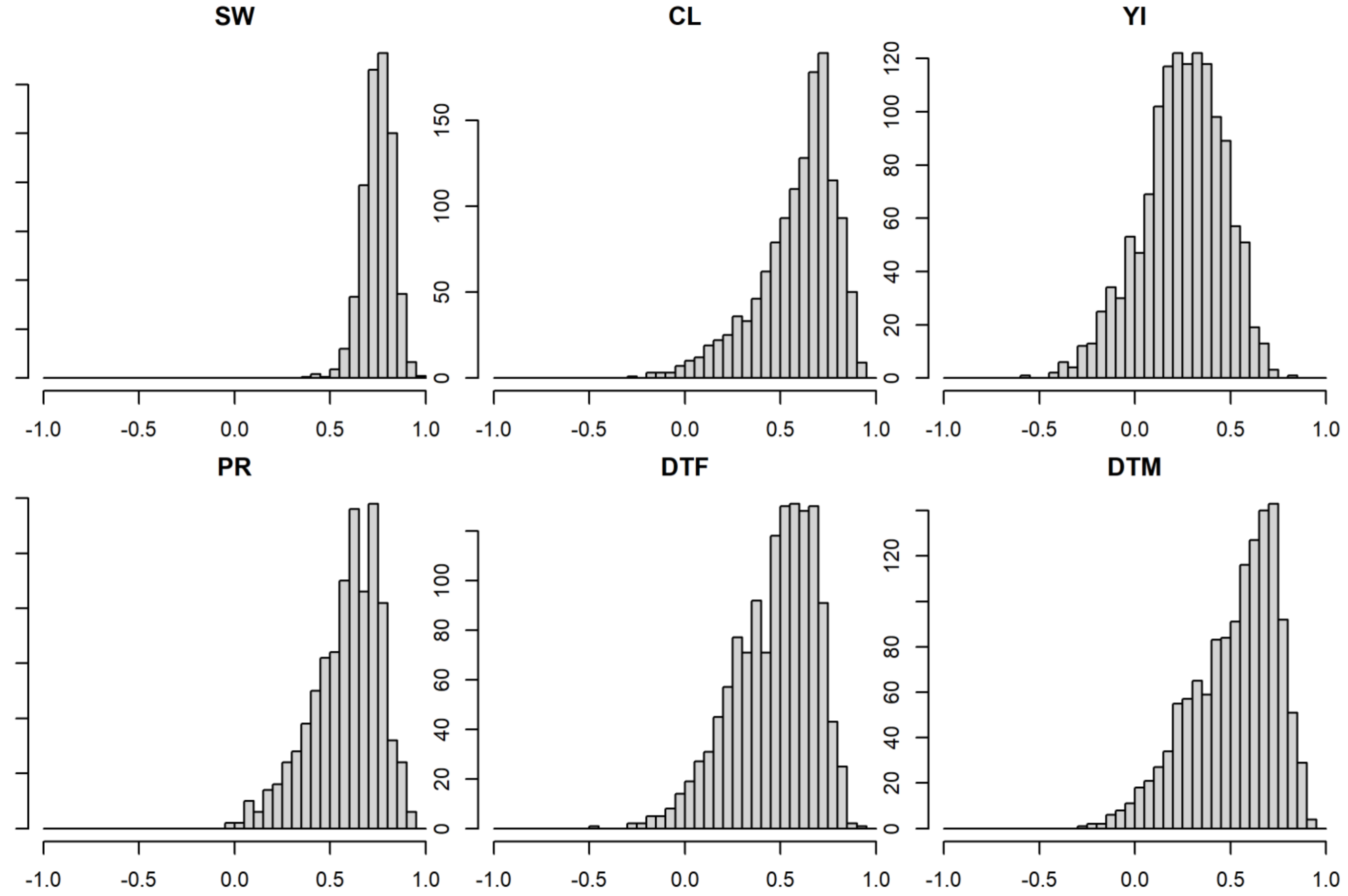


**Figure S3 Distributions of estimates of genetic correlations between environments**

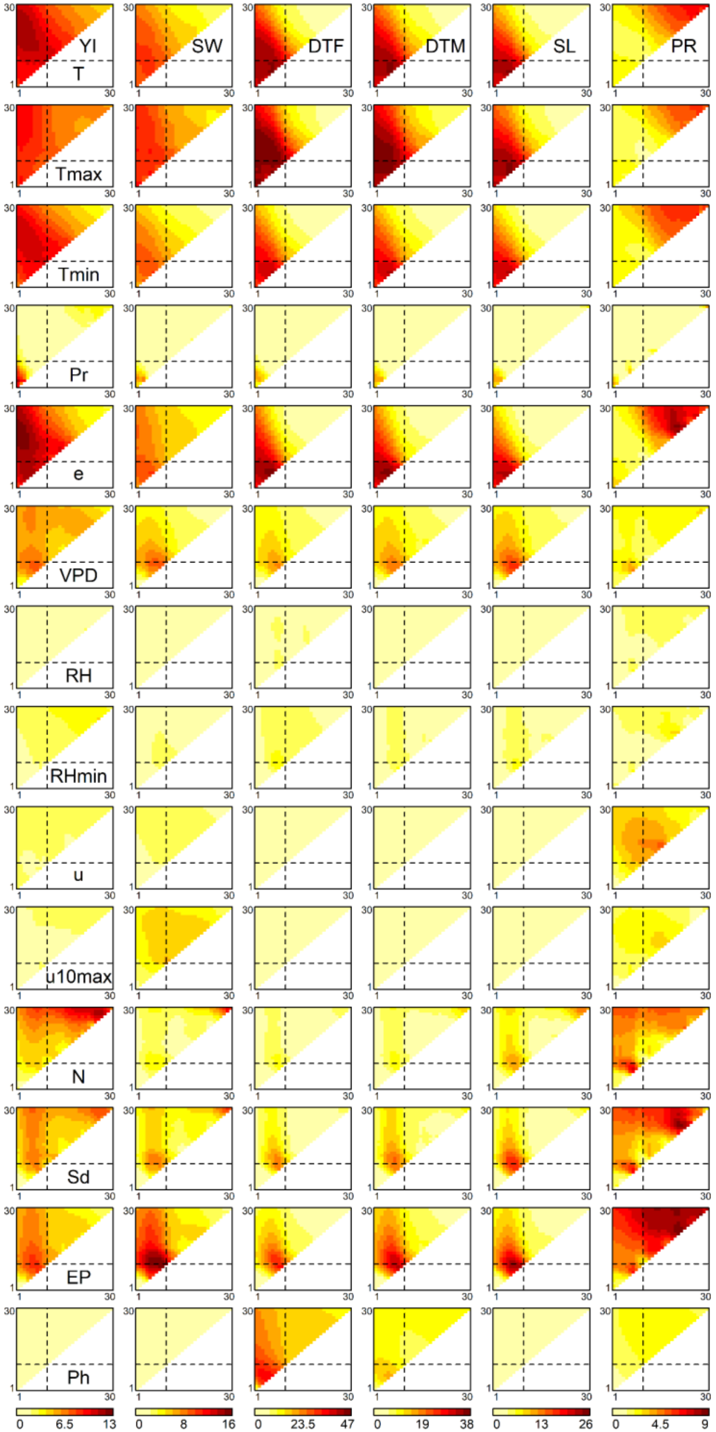


**Figure S4 Associations of environmental covariates with genetic correlations between environments.** The heat maps represent the −log10 *P* values for the correlation coefficients ($r^2$) of off-diagonal elements between the similarity matrix of each environmental covariate and the genetic correlation matrix. The diagonal boxes of the triangles correspond to the 1st to 30th growth stages, from the lower left to the upper right. The off-diagonal elements correspond to the growth periods that span multiple stages, where the x and y axes denote the start and end of the periods, respectively. The broken lines indicate flowering time.

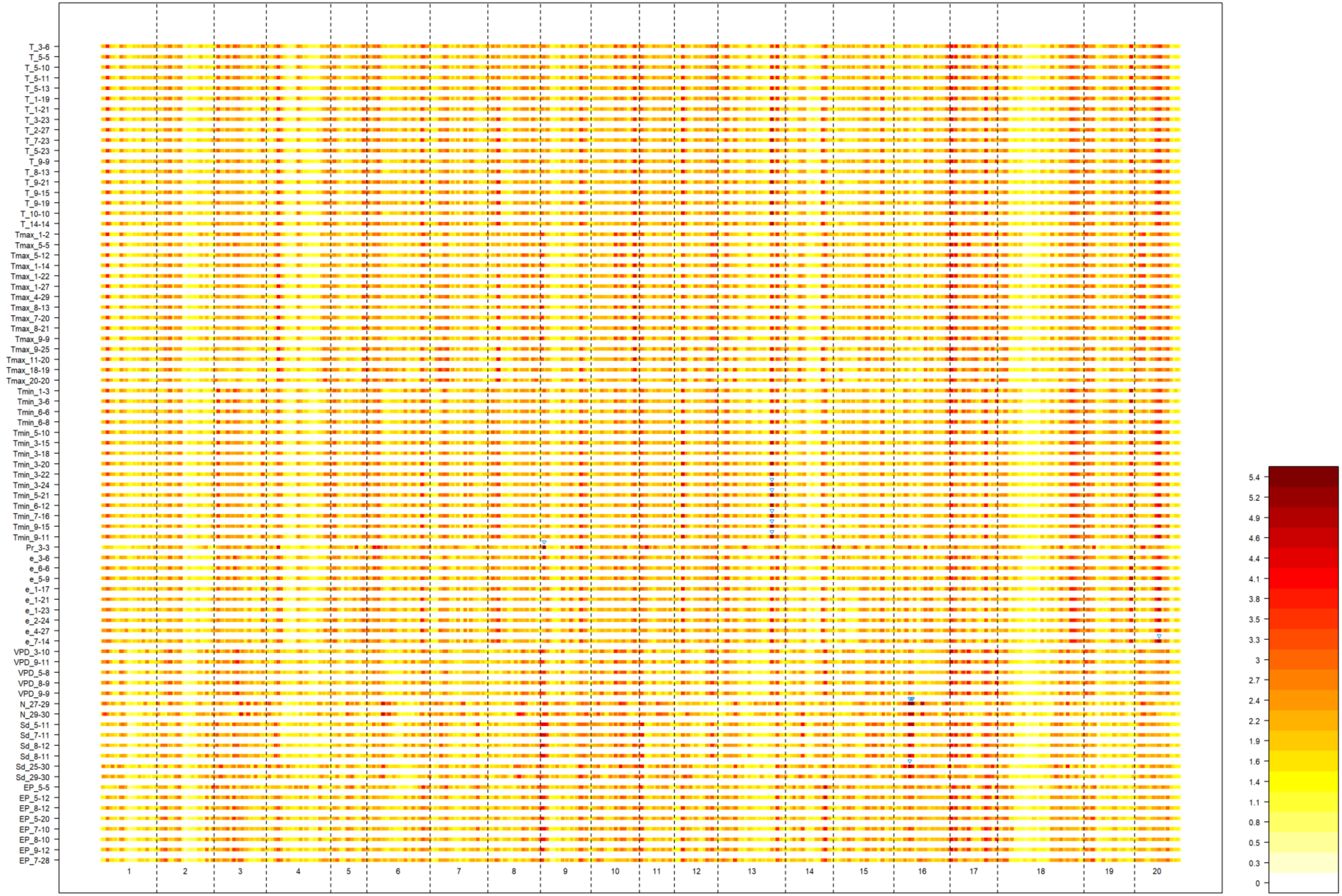


**Figure S5 Results of the genome-wide association mapping for the slopes obtained for seed weight.** The x and y axes indicate the chromosomal positions and the environmental covariates, respectively. The −log10 *P* values are illustrated using colours. The blue triangles indicate significant SNPs with a false discovery rate <0.05.

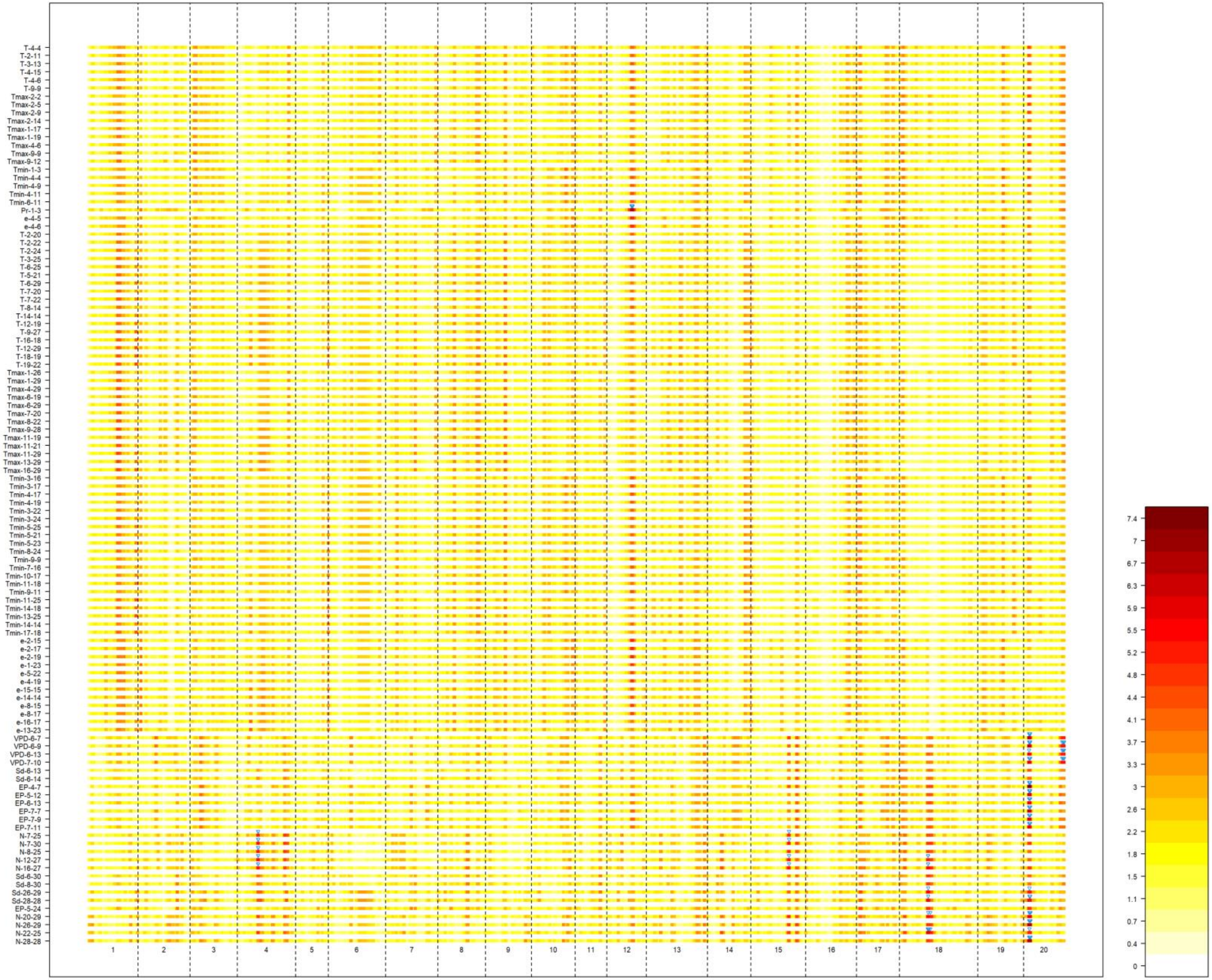


**Figure S6 Results of the genome-wide association mapping for the slopes obtained for yield.** The x and y axes indicate the chromosomal positions and the environmental covariates, respectively. The −log10 *P* values are illustrated using colours. The blue triangles indicate significant SNPs with a false discovery rate <0.05.

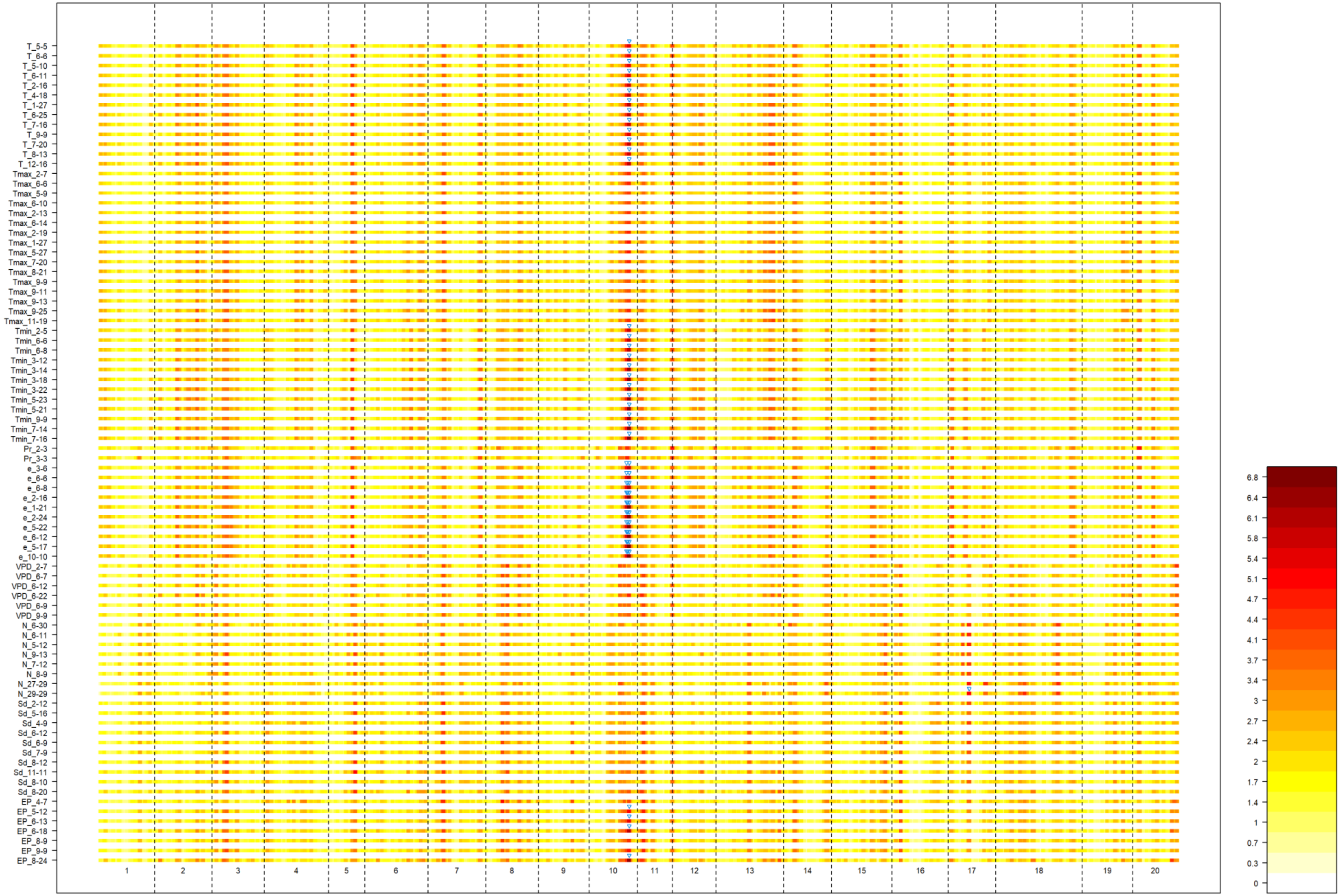


**Figure S7 Results of the genome-wide association mapping for the slopes obtained for stem length.** The x and y axes indicate the chromosomal positions and the environmental covariates, respectively. The −log10 *P* values are illustrated using colours. The blue triangles indicate significant SNPs with a false discovery rate <0.05.

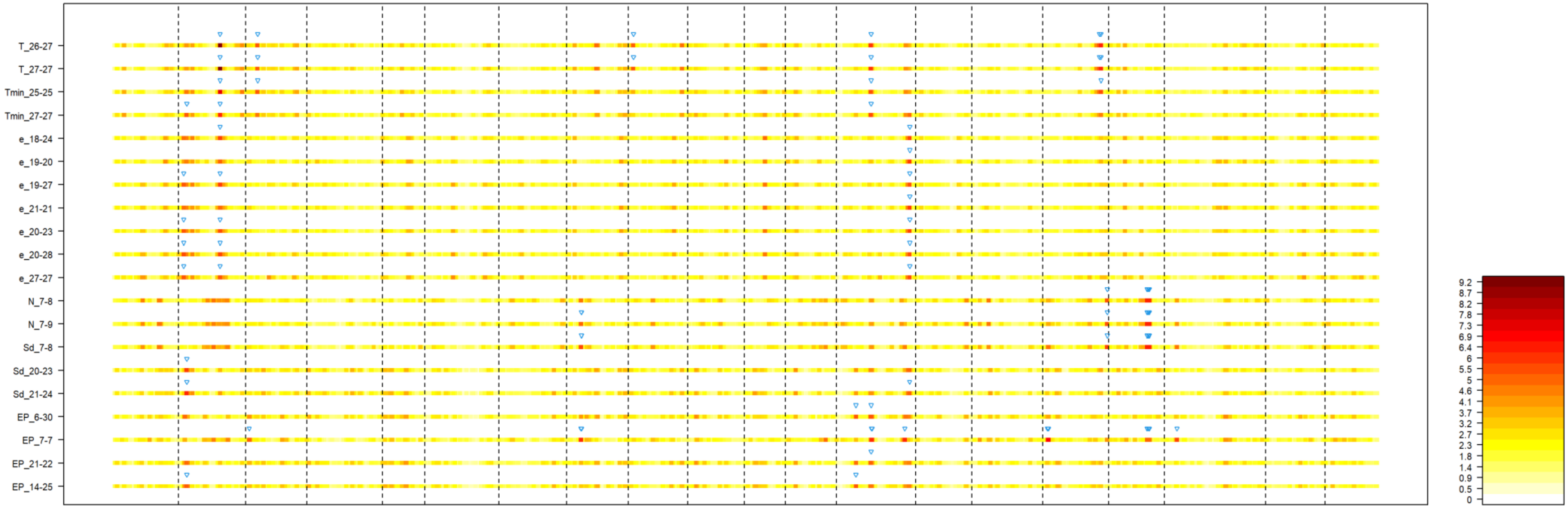


**Figure S8 Results of the genome-wide association mapping for the slopes obtained for protein content.** The x and y axes indicate the chromosomal positions and the environmental covariates, respectively. The −log10 *P* values are illustrated using colours. The blue triangles indicate significant SNPs with a false discovery rate <0.05.

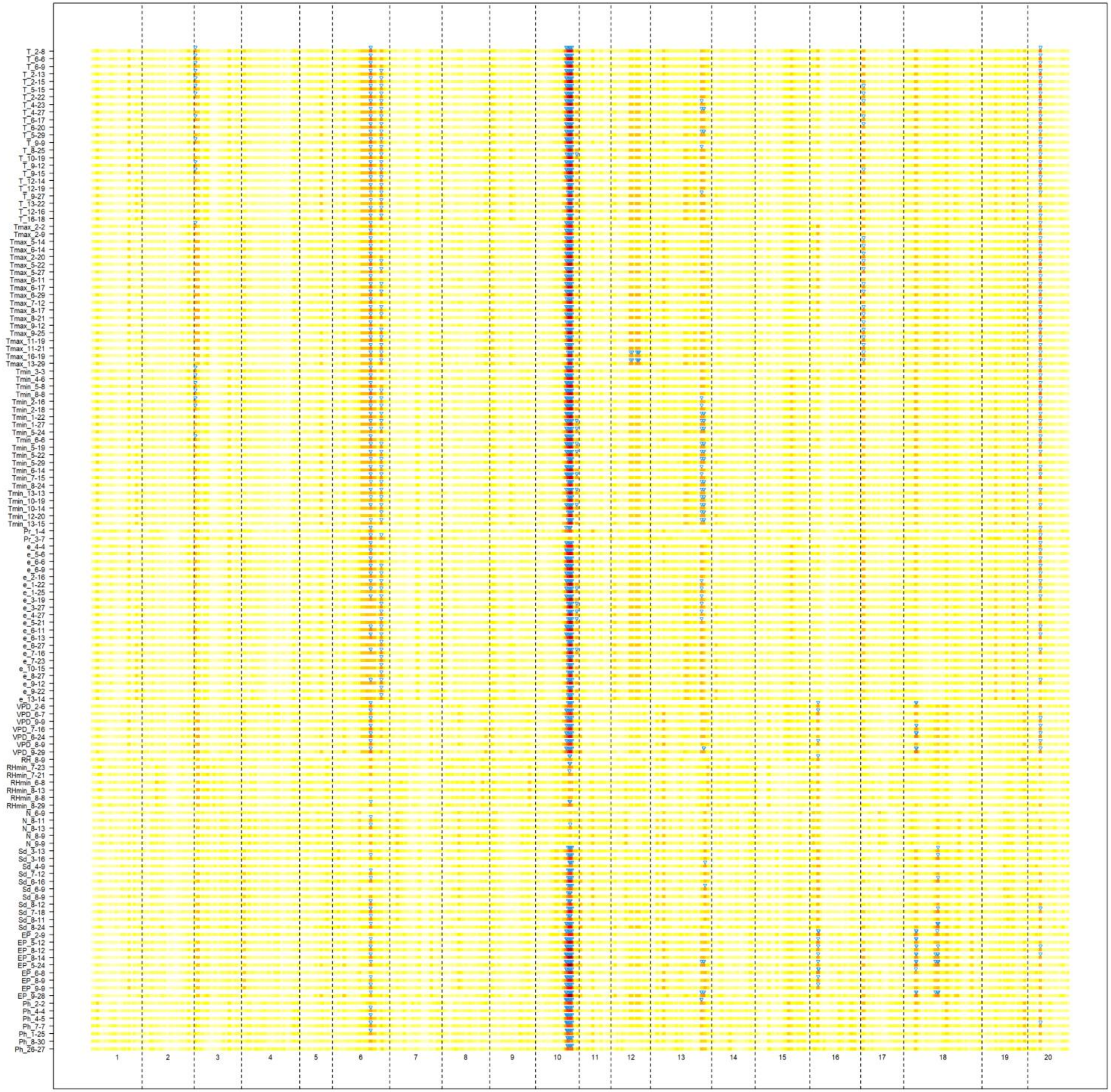


**Figure S9 Results of the genome-wide association mapping for the slopes obtained for days to flowering.** The x and y axes indicate the chromosomal positions and the environmental covariates, respectively. The −log10 *P* values are illustrated using colours. The blue triangles indicate significant SNPs with a false discovery rate <0.05.

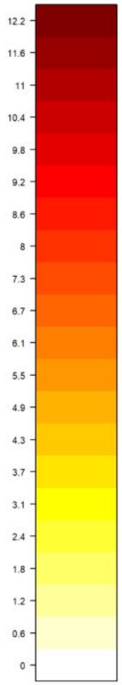

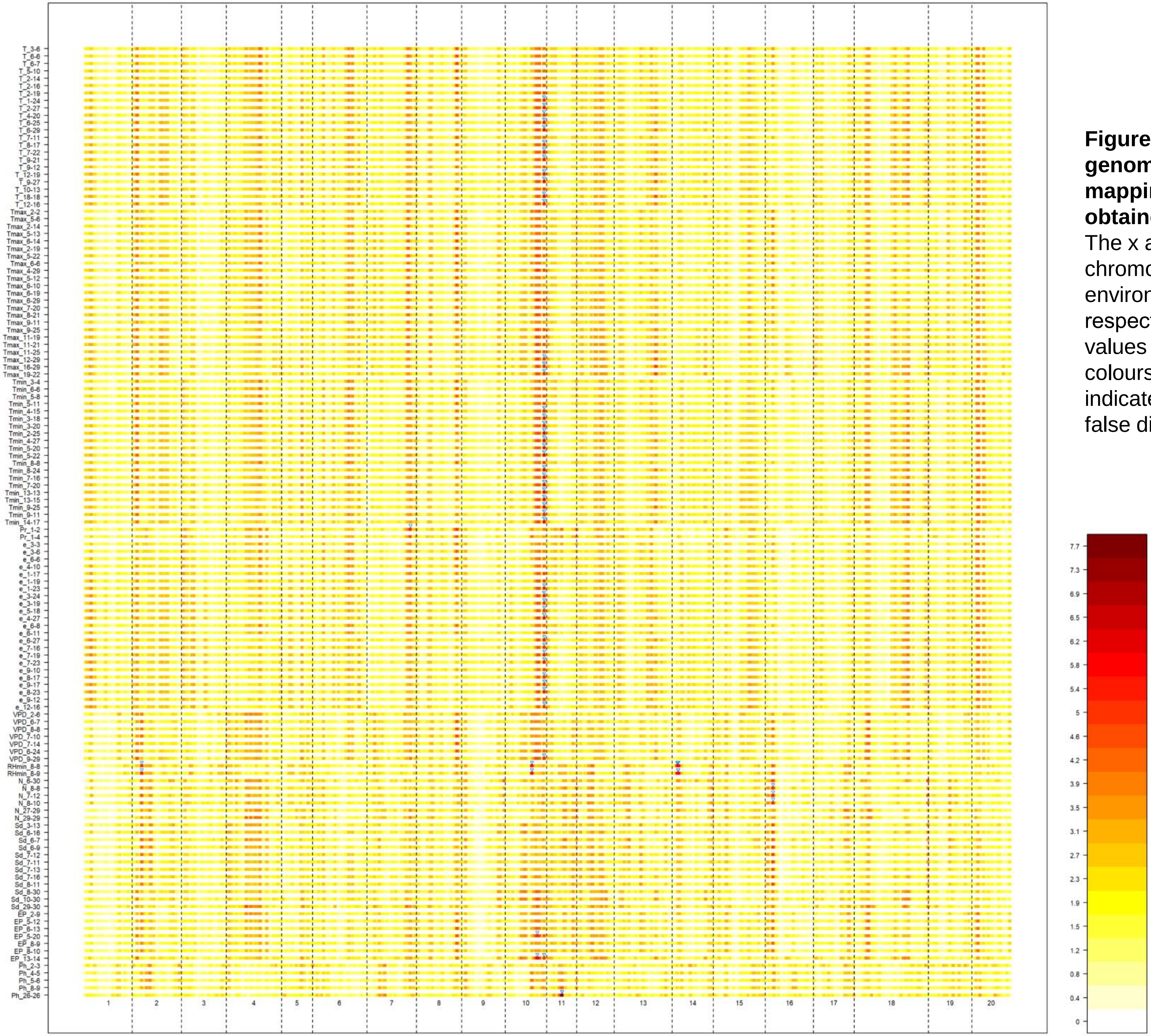


**Figure S10 Results of the genome-wide association mapping for the slopes obtained for days to maturity**. The x and y axes indicate the chromosomal positions and the environmental covariates, respectively. The −log10 *P* values are illustrated using colours. The blue triangles indicate significant SNPs with a false discovery rate <0.05.

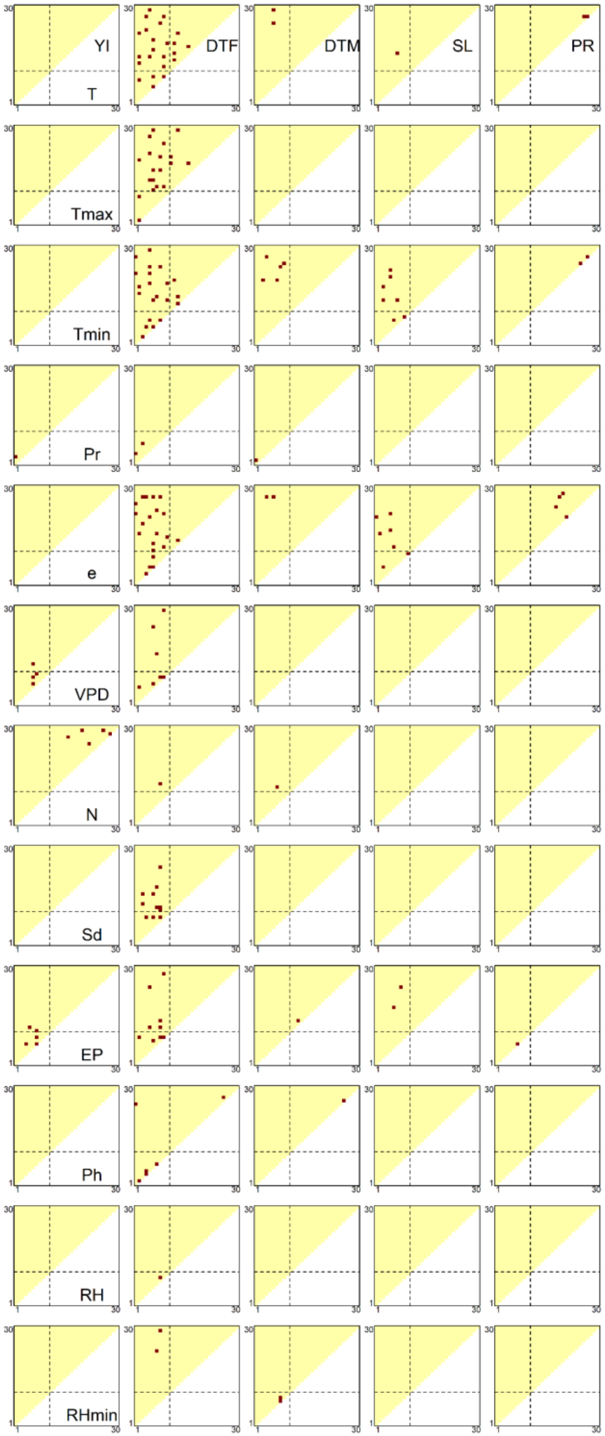


**Figure S11 Distributions of meteorological factor/growth stage combinations where significant associations were detected.** Red dots indicate the combinations with significant associations. The diagonal elements of the triangles correspond with the 1st to 30th growth stages, from the lower left to the upper right. The off-diagonal elements correspond to the growth periods that span multiple stages, where the x and y axes denote the start and end of the periods, respectively.

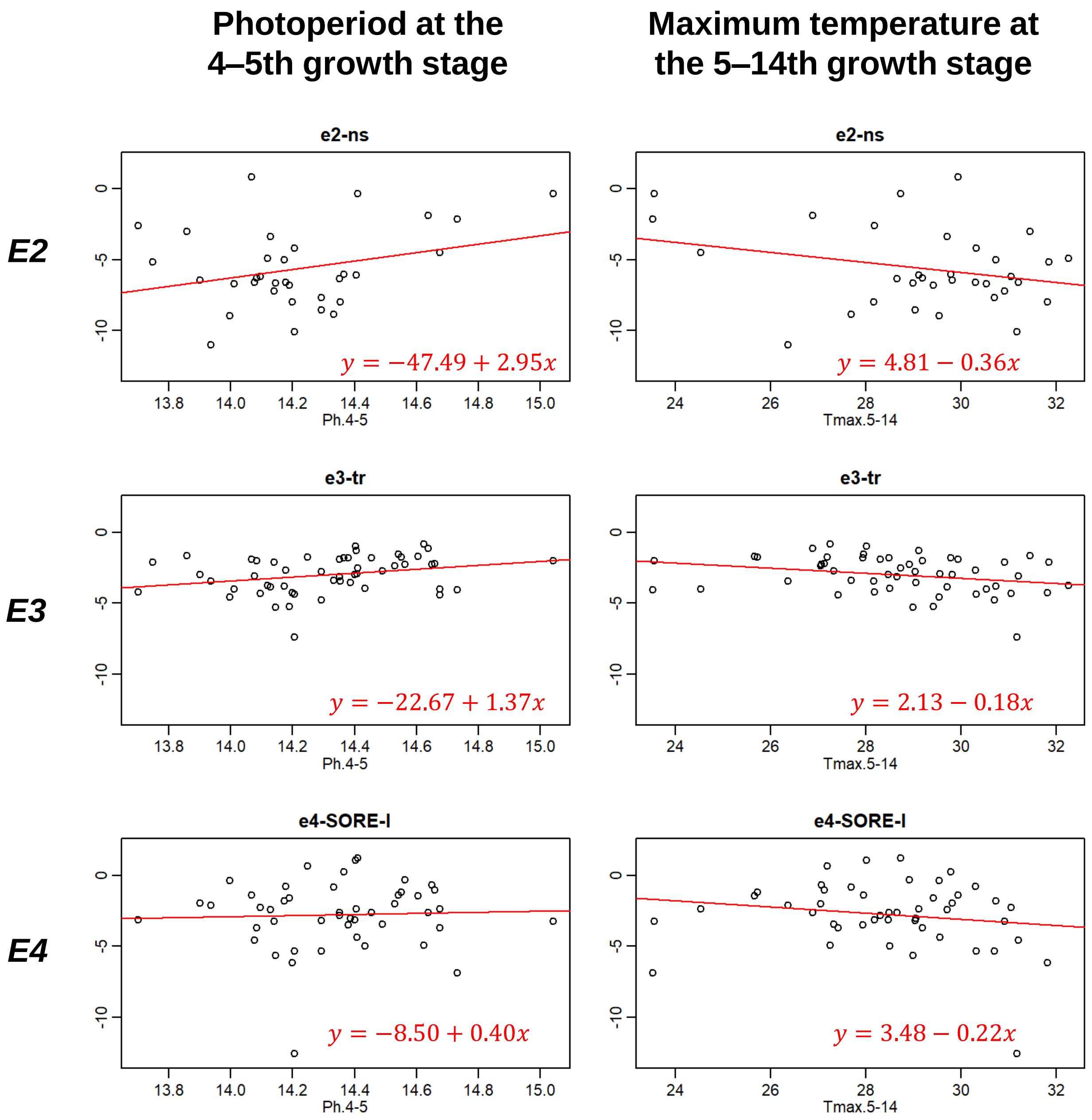


**Figure S12 Allele substitution effects of flowering genes (*E2*, *E3* and *E4*) on DTF.** The effects of loss-of-function alleles (*e2-ns*, *e3-tr* and *e4-SORE-l*) were estimated for each environment (Supplementary Methods) and plotted against two environmental covariates (photoperiod at the 4$^{th}$ to 5$^{th}$ growth stages and maximum temperature at the 5$^{th}$ to 14$^{th}$ growth stages). For both environmental covariates, *E2* showed the greatest slopes, suggesting that *E2* can affect the G × E interactions of DTF. The red lines are the regression lines estimated using the least squares method.
